# Defining and cataloging variants in pangenome graphs

**DOI:** 10.1101/2025.08.04.668502

**Authors:** Pouria Salehi Nowbandegani, Shenghan Zhang, Haoyang Hu, Heng Li, Luke J. O’Connor

**Affiliations:** Department of Biomedical Informatics, Harvard Medical School, Boston, MA, USA; Program in Medical and Population Genetics, Broad Institute of MIT and Harvard, Cambridge, MA, USA; Department of Data Science, Dana-Farber Cancer Institute, Boston, MA, USA

## Abstract

Structural variation causes some human haplotypes to align poorly with the linear reference genome, leading to ‘reference bias’. A pangenome reference graph could ameliorate this bias by relating a sample to multiple reference assemblies. However, this approach requires a new definition of a ‘genetic variant.’ We introduce a definition of pangenome variants and a method, *pantree*, to identify them. Our approach involves a pangenome *reference tree* which includes all nodes (sequences) of the pangenome graph, but only a subset of its edges; non-reference edges are *variant edges*. Our variants are biallelic and have well-defined positions. Analyzing the Minigraph-Cactus draft human pangenome reference graph, we identified 29.6 million genetic variants. Most variants (99.2%) are small, and most small variants (73.9%) are SNPs. 3.5 million variants (11.7%) have a reference allele which is not on GRCh38; these variants are difficult to detect without a pangenome reference, or with existing pangenome-based approaches. They tend to be embedded within tangled, multiallelic regions. We analyze two medically relevant regions, around the HLA-A and RHD genes, identifying thousands of small variants embedded within several large insertions, deletions, and inversions. We release an open-source software tool together with a VCF variant catalogue.

## Introduction

Human genomes vary in myriad ways, from short polymorphic repeats to large scale rearrangements. In regions with structural variation, many haplotypes align poorly with any single reference genome; two haplotypes may be similar to each other but dissimilar to the reference, for example when they carry different alleles of a long insertion. A natural solution is to collate multiple reference assemblies, comprising a *pangenome*, and in 2023, the Human Pangenome Reference Consortium (HPRC) published a draft pangenome reference including 47 diploid assemblies (Eizenga et al., 2020; Garrison et al., 2018; Hickey et al., 2024; Liao et al., 2023; Sedlazeck et al., 2018). The pangenome reference requires a change in perspective. A single genome is represented not as a linear sequence of alleles but rather as a walk through the pangenome graph, in which each node represents a sequence of base pairs. The notion of genomic ‘position’ must be revisited, as many sequences are missing from the linear reference. Most fundamentally, it becomes a challenge to identify or even to define a genetic ‘variant’.

Current approaches define variants as different types of ‘bubbles,’ which are regions of the graph where two walks might split apart and merge back together. Specifically, the definition adopted by HPRC was that a variant is a *superbubble*, flanked by a single entry node and a single exit node at which every genome aligns (Onodera et al., 2013). In regions harboring large-scale structural variation, superbubbles may contain numerous long alleles, some of them differing by just a few nucleotides despite being kilobases in length. One proposed solution is to align these alleles back to the reference genome; this approach recovers small variants nested within deletions, but not insertions or other non-reference sequences (Liao et al., 2023). Another is to search for superbubbles within other superbubbles hierarchically; this approach succeeds when variants are nested inside each other, but not when they are mutually interlocking, and the same genetic difference can reappear at multiple levels of the hierarchy (Paten et al., 2018). More generally, these approaches fall short of addressing the conceptual problem. What is a pangenome variant definitionally?

We propose to define pangenome variants against a *reference tree*. This tree contains all the nodes of the pangenome graph, but only a subset of its edges. The remaining edges, called *variant edges*, represent genetic variants: when some genome passes through a variant edge, it carries an alternative allele; when it reaches the same node via a reference tree edge, it carries a reference allele. With this definition, alleles are paths on the reference tree, and variants are pairs of alleles with a shared ‘branch point’. Mathematically, the set of variant edges is akin to a basis for a vector space.

This definition satisfies our intuition, it has the right mathematical properties, and it solves various practical problems. We define a reference tree for the HPRC draft pangenome and produce a variant catalog including many thousands of non-GRCh38 variants. We compare our variants with previously identified superbubbles, characterizing the differences between different approaches. We analyze two complex, medically relevant loci, visualizing and cataloguing the variation that is present in these regions. We make our variant catalog available as a variant call format (VCF) file, which is compatible with widely used tools.

## Results

### Definition of a variant

In a pangenome graph, each node represents a DNA sequence. A sample is represented as a walk, the nodes of which are concatenated. Many parts of the graph have a simple linear structure, forming a sequence of separate superbubbles. Other parts contain nested structure, for example, when a SNP is embedded inside an insertion, or interlocking structure, for example, when two different deletions partially overlap. Some regions can be particularly complex, especially in tandem repeats and segmental duplications.

Figure 1a shows a possible pangenome graph. The blue subgraph in Figure 1b is a reference tree, and the remaining edges, in red, are variant edges. The genotype of a walk is defined as the number of times that it visits each variant edge; the green walk (Figure 1c) has genotype [0,1,1] for variant edges (1,4), (7,8), (9,3) respectively. In a tree, there exists a unique path between any two nodes. Thus, a walk that traverses exactly one variant edge (*u, v*) must follow the unique reference tree path from its starting node to *u*, and then the unique reference path from v to its ending node. More generally, a walk can be reconstructed from its genotype (see Supplementary Note, Section 2, Theorem 2.1).

**Figure 1:**
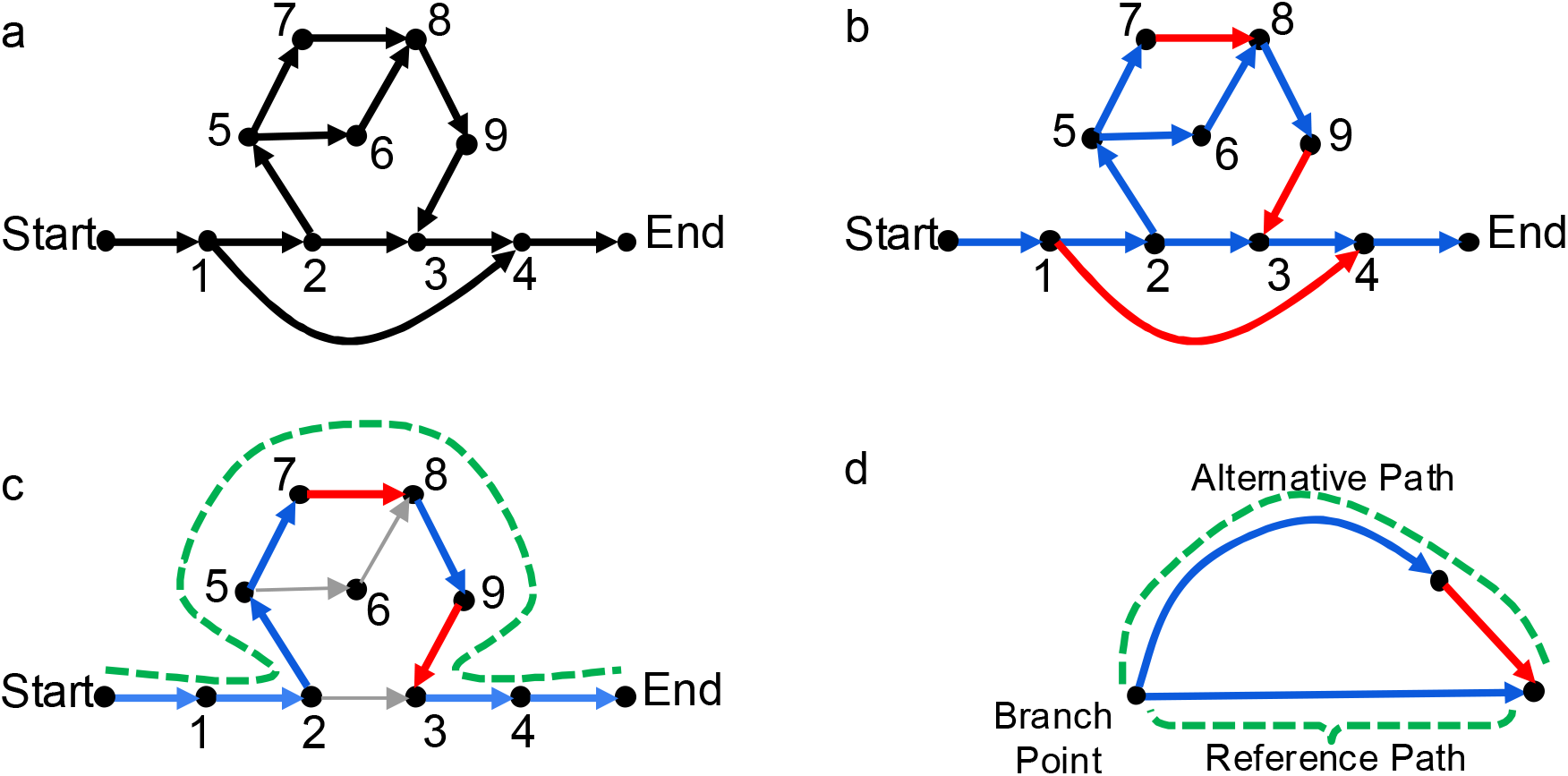
Definition of a pangenome variant. (a) A toy example graph with 11 nodes. (b) A reference tree (in blue) and corresponding variant edges (in red). (c) A walk representing one individual haplotype (in green), with its variant edges highlighted (red). (d) Paths corresponding to the reference and alternative alleles of a variant edge.

A set of variant edges gives a basis for a vector space, where the elements of that space are differences between walks (see Supplementary Note, Section 6). Different choices of reference tree yield different variant edges and different reference vs. alternative alleles, but any choice gives a complete description of the ways in which individual genomes differ. Moreover, some features are invariant. Node 3 in Figure 1a has two incoming edges, one of which will be chosen as the reference tree edge; the other will be a variant edge.

Because DNA is a double-stranded molecule, pangenome graphs are *bidirected*, respecting the symmetry between a sequence and its reverse complement (e.g., ATCC and GGAT) (MYERS, 1995). In the Supplementary Note, we provide precise definitions for a bidirected graph, a bidirected spanning tree, and a bidirected variant edge. A choice of spanning tree naturally orients each node of a bidirected graph; variant edges between nodes of different orientation are *inversion edges*. A simple inversion event corresponds to two different inversion edges: one deletes a DNA sequence, and the other inserts its reverse complement.

Many bioinformatics pipelines – in particular, those based on the VCF file format – require variants to have well-defined positions with respect to a linear reference. We include the linear reference (here, GRCh38) as a branch of the reference tree and define the position of a non-GRCh38 node u as the highest position along the linear reference such that the reference tree contains a directed path from that position to *u*. A variant edge (*u, v*) has position (*pos*(*u*), *pos*(*v*)), although this definition is modified for the VCF (see Methods and Discussion). This definition allows us to work locally with respect to the linear reference: within a region of interest, it suffices to consider just those variant edges whose position overlaps with the region (see Supplementary Note, Section 4, Theorem 4.1). The same is not true if variant position is defined as any single point. We also use variant positions to define haplotype missingness (see Supplementary Note, Section 10).

Variant edges have a reference allele and an alternative allele, which are defined with respect to the *branch point* of a variant edge. The branch point of non-inversion variant edge (*u, v*) is the lowest common ancestor of *u* and *v* in the reference tree (Figure 1d). The reference allele corresponds to the reference tree path from the branch point to *v*, excluding *v*; the alternative allele corresponds to the reference tree path from the branch point to *u*, including *u* (see Supplementary Note, Section 7).

We classify variant edges into five non-overlapping categories: insertions, deletions, replacements, duplications, and inversions (see Supplementary Note. Section 8). An edge that skips forward from an ancestor to a descendant in the reference tree is a deletion. An edge that returns backward from a descendant to an ancestor is a duplication. An edge that crosses between two different branches is either an insertion, if its reference allele is empty, or a replacement; single nucleotide polymorphisms (SNPs) are replacements whose alleles both have length one, and multiple nucleotide polymorphisms (MNPs) are replacements whose alleles have the same non-unit length. An inversion is an edge connecting two nodes with different orientations, and they normally occur in pairs. For example, in Figure 1b, the variant edge (1,4) is a forward edge representing a deletion; the edge (7,8) is a crossing edge representing a replacement; and the edge (9,3) is a crossing edge representing an insertion.

### Catalogue of variants

We analyzed the draft human pangenome reference compiled by HPRC, in particular the genome graph inferred using Minigraph-Cactus (MC) (Hickey et al., 2024) (see Data Availability). This graph contains 80,069,733 nodes, 110,938,345 edges, and 90 haplotypes, including the GRCh38 reference assembly. Each haplotype is represented by 113-566 contiguous walks, with an average of 286 walks per haplotype. Most samples in the reference panel are from the 1000 Genomes project and have African or American ancestry. We applied our software package (see Methods and Code Availability) and produced a single VCF file containing the variants that we found, together with the genotype of each sample (see Methods and Data Availability).

We identified 29.6 million genetic variants and annotated them as either a SNP, an insertion, a deletion, an MNP, a replacement (non-SNP and non-MNP), a duplication, or an inversion. Variants with a combined allele length of <50bp were called ‘small’, others ‘large’. As expected, most variants (99.2%) were small, and most small variants (73.9%) were SNPs, with a substantial number of small insertions (10.8%) and deletions (12.1%) (**Figure 2a)**. Most large variants were insertions (38.0%) or deletions (41.3%) (**Figure 2b**). Most small indels were repeat expansions or contractions (**Figure 2a**), most often short tandem repeats and homopolymers (**Supplementary Figure 1**). Because of the way that Minigraph-Cactus constructs the graph, most repeat expansions are annotated as insertions, and not duplications; there were just 53 duplications. The alternative PGGB algorithm (Garrison et al., 2024) is more likely to produce a duplication. There were 860 inversion edges, almost all of them large, and these inversion edges correspond to about half as many inversion events (see Discussion). These numbers include chromosomes 1-22; chromosomes X and Y had similar distributions of variant types (**Supplementary Figure 2 and Supplementary Figure 3**).

**Figure 2:**
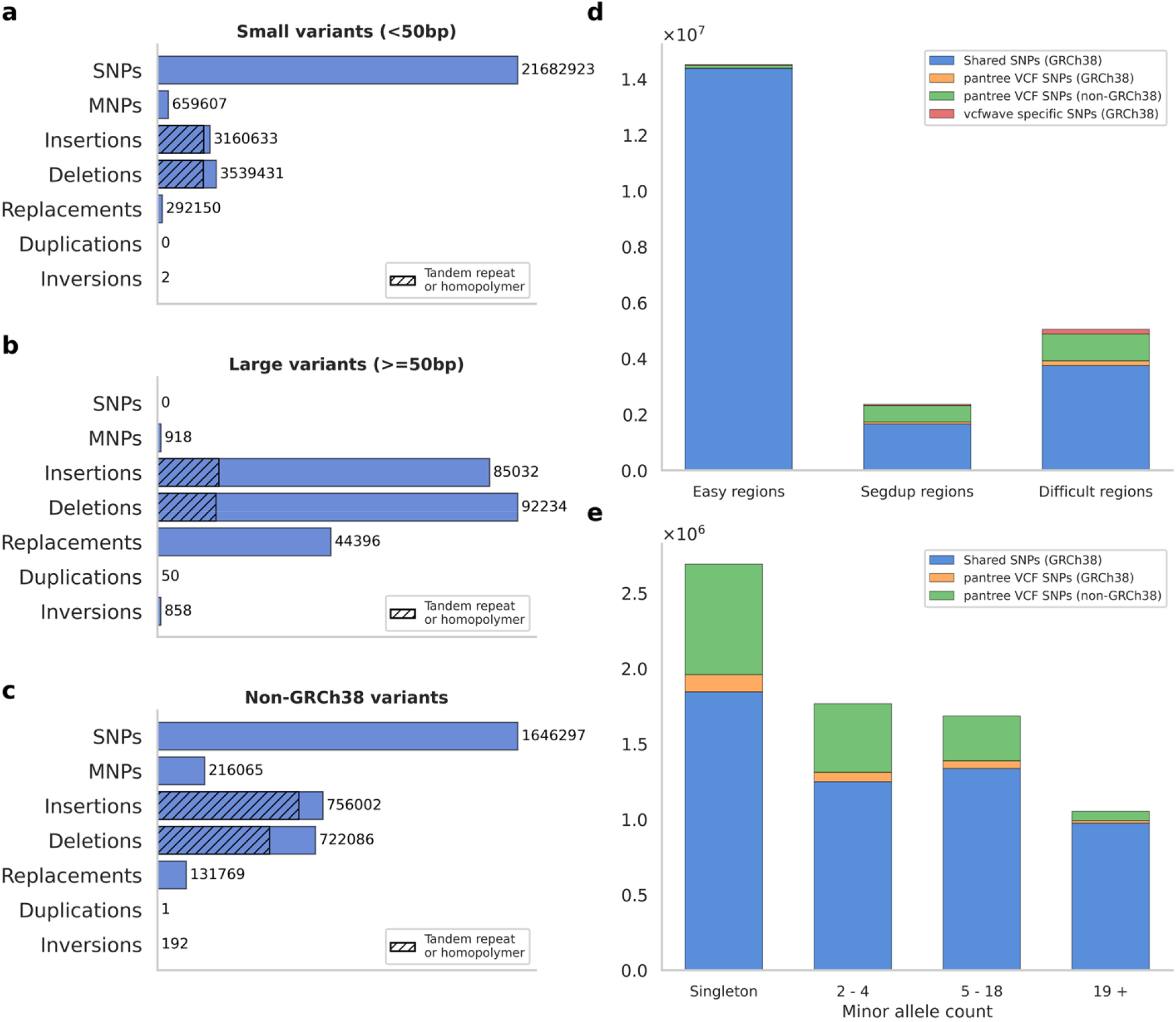
Summary of variants in the HPRC draft pangenome. (a) Number of small variants (<50bp combined allele length) across seven non-overlapping variant types. Hatched area indicates the number of variants that cause a repeat expansion or contraction. (b) Number of large variants (>=50bp). (c) Number of non-GRCh38 variants, whose reference allele is not on the GRCh38 genome. (d) Overlap of SNPs with those reported by *vcfwave*, across three non-overlapping genome annotations. (e) Minor allele count (the minimum of the reference and alternative allele counts) out of 90, stratified by overlap with *vcfwave*. For numerical results, see **Supplementary Tables 1-5**.

3.5 million variants (11.7%) had a reference allele that is not on the linear reference genome, and we refer to these as non-GRCh38 variants. About half were SNPs, others mostly indels (**Figure 2c**). These variants are difficult to detect without a pangenome reference. They include 21,109 variants, mostly indels, that are both common and large (**Supplementary Figure 4**).

We compared the SNPs that were detected using our approach vs. the ‘superbubble’ approach that was previously used by HPRC (see Methods). Superbubbles are disjoint subgraphs separated from the rest of the graph by pinch-point nodes - one ‘entrance’ and one ‘exit’ - where every haplotype aligns, including the linear reference (Paten et al., 2018). In **Figure 1a**, the entire graph is a single superbubble. In ‘easy’ regions of the genome, where every haplotype closely tracks the linear reference genome, both approaches found approximately the same set of SNPs. In segmental duplications, our approach detected 46.1% more SNPs (2,313,929 vs. 1,584,132), and in other difficult regions, it detected 43.4% more SNPs (**Supplementary Figure 5**). The SNPs that we find are a superset of those that are superbubbles except for exactly two superbubble- specific SNPs (these arise due to assembly gaps; (**Supplementary Figure 6 and Supplementary Figures 7**). All non-superbubble SNPs are nested within larger superbubbles. 62.0% of non-superbubble SNPs are non-GRCh38 SNPs.

For some non-superbubble SNPs, an existing approach, *vcfwave* (Liao et al., 2023), can recover them by re-aligning the alternative alleles of a superbubble with its reference allele. This approach cannot recover non-GRCh38 SNPs, as it relies on pairwise alignment with the linear reference genome (see Methods). We find 1,896,587 SNPs that are missed by *vcfwave*, most of which are non-GRCh38 **(Figure 2d)**. Non-vcfwave SNPs are modestly enriched at lower allele frequencies, but many of them (23.8%) are common (minor allele count >= 5 out of 90) (**Figure 2e**). vcfwave variants can be inconsistent with the pangenome graph, and indeed, 239,375 SNPs are vcfwave-specific (**Figure 2d**); these arise because *vcfwave* may produce a different alignment compared with the graph construction algorithm (**Supplementary Figure 8**). Chromosome X had a similar proportion of non-GRCh38 SNPs and non-vcfwave SNPs as compared with the autosomes (**Supplementary Figure 2**); chromosome Y was enriched for non-GRCh38 and non-vcfwave SNPs (**Supplementary Figure 3**).

Minigraph-Cactus uses a linear reference genome, here GRCh38, to initialize graph construction. Regions that are missing from GRCh38 entirely, including centromeres and acrocentric arms, are also missing from the Minigraph-Cactus GRCh38 graph. We analyzed the HPRC Minigraph-Cactus CHM13 graph, which is initialized at the telomere-to-telomere CHM13 assembly (see Data Availability). This graph contains 14.7% more variants, and the difference is mostly explained by an increased number of variants within centromeres, subtelomeres, and acrocentric arms (**Supplementary Figure 9a**). These regions are highly enriched for off-linear-reference variants; the number of non-CHM13 variants in the CHM13 graph is 42.4% greater than the number of non-GRCh38 variants in the GRCh38 graph, almost entirely because of these regions (**Supplementary Figure 9b**). Graph construction in these regions is difficult, and some of these variants could be alignment errors. In the GRCh38 graph, most non-GRCh38 variants are not contained in these regions.

### Multiallelic superbubbles

All non-GRCh38 variant edges are contained within a superbubble containing at least two variant edges (**Figure 3a**). These multi-variant superbubbles are usually multiallelic; we sought to characterize multiallelic, multi-variant superbubbles. Both count distributions -the number of alleles per superbubble, and the number of variant edges -were heavy-tailed (**Figure 3b**). There were 23,694 superbubbles containing at least 20 alternative alleles, and these contained 11.1% of variant edges (**Figure 3c**). Allele count and variant count are correlated, but there is no exact correspondence (**Figure 3d**), except that superbubbles with more than one alternative allele always contain more than one variant edge.

**Figure 3:**
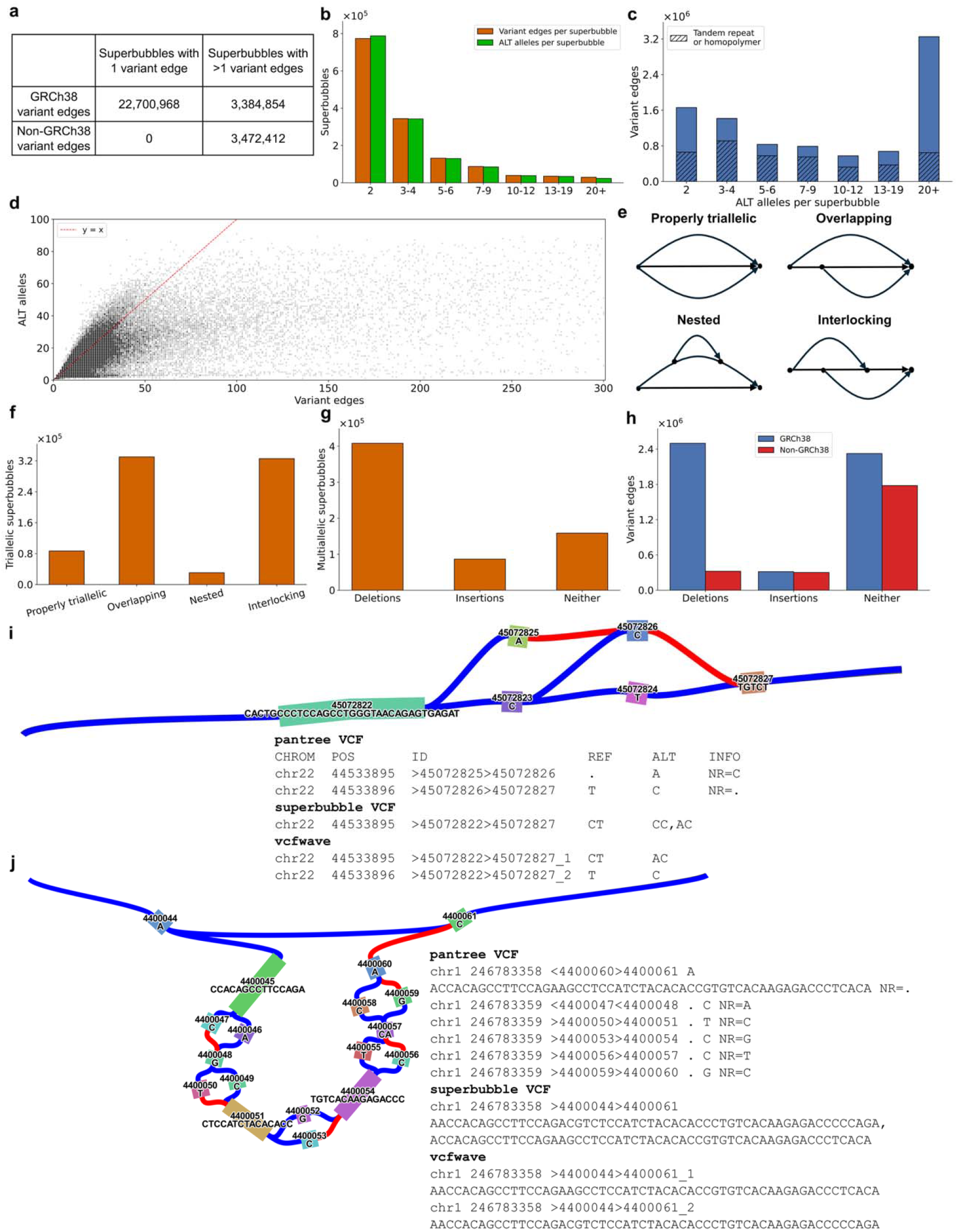
Variant content of multi-allelic superbubbles. (a) Number of GRCh38 and non-GRCh38 variant edges contained in biallelic (1 variant edge) versus multiallelic (>1 variant edge) superbubbles. All non-GRCh38 variant edges occur in multiallelic superbubbles. (b) The number of multiallelic superbubbles containing different numbers of variant edges (orange) and alternative alleles (green). (c) The number of variant edges contained in superbubbles with different numbers of alternative alleles. Hatched bars indicate the number of repeat expansions or contractions. (d) Number of alternative alleles vs. variant edges per superbubble. (e) Schematic of each possible triallelic superbubble. (f) Counts of each possible triallelic superbubble. (g) Number of more-than-triallelic superbubbles classified as insertions, deletions, or neither. (h) Number of GRCh38 and non-GRCh38 variants in more-than-triallelic superbubbles classified as insertions, deletions, or neither. (i) Example of a triallelic superbubble, with excerpts from three VCF files. The columns are chromosome, position, node identifiers, reference allele, and alternative allele or alleles, respectively. (j) Example of a multi-allelic superbubble classified as an insertion, with VCF excerpts. For numerical results, see **Supplementary Tables 6-10**.

We classified multiallelic superbubbles in three ways. First, we classified triallelic, two-variant superbubbles as either ‘properly triallelic’, ‘overlapping’, ‘nested’, or ‘interlocking’ (**Figure 3e**) (see Methods). Most triallelic superbubbles are either interlocking (42.1%) or overlapping (42.7%) (**Figure 3f**). Variants in these bubbles are mostly SNPs (47.0%) or indels (48.4%) (**Supplementary Table 9**), and they account for 3.5% of non-GRCh38 variants. The predominant ‘interlocking’ and ‘overlapping’ categories are difficult to decompose using existing approaches. Properly triallelic superbubbles do not need to be decomposed, as their three alleles have no partial similarity. Nested superbubbles can be decomposed using an existing hierarchical approach, which is to remove the endpoints of a superbubble and identify superbubbles within the connected components of the remaining subgraph, but this approach cannot decompose overlapping or interlocking superbubbles. *vcfwave* sometimes resolves overlapping or interlocking variants, but its behavior depends on which of the three alleles is the reference allele; for example, when two SNPs are interlocking and one of them is off of the linear reference, *vcfwave* does find two variants, but only one of them is a SNP (**Figure 3i**). It also misses a handful of GRCh38 SNPs due to a rare edge case (**Supplementary Figure 10**).

Second, we classified multiallelic superbubbles with more than three alleles as insertions (13.2%), deletions (62.5%), or neither (24.3%) (**Figure 3g**). An insertion superbubble contains a reference tree edge from the beginning to the end of the superbubble (**Figure 3j**); a deletion superbubble contains a beginning-to-end variant edge. The larger number of deletions is mostly due to an ascertainment effect, that deletions are more likely to be multiallelic (**Supplementary Figure 11**). Most non-GRCh38 variants (74.0%) are found outside of insertion and deletion superbubbles, although insertions (which comprise non-GRCh38 sequence exclusively) are enriched, containing 12.6% of non-GRCh38 variants in total (**Figure 3h**).

Third, in order to understand where non-GRCh38 variants are located, we focused on highly multiallelic superbubbles, with at least ten alternative alleles. There are only 95,632 such bubbles (**Figure 3b**), but each one may contain dozens of variants (**Figure 3d**). These contain 65.9% of non-GRCh38 variants. They are likely to contain long insertions without being classified strictly as an insertion superbubble; 2.1% of them contain an insertion of at least 1kb, and these contain 37.6% of non-GRCh38 variants, an average of 646 per superbubble.

### Complex regions

We investigated two specific regions of the genome which are known to harbor complex patterns of genetic variation: the region around the HLA-A gene, which contains HLA-A and several copy number-variable pseudogenes, and the RHD region, which contains an inversion overlapping the RHD gene (see Methods). HLA alleles have well-studied effects on autoimmunity, and RHD alleles largely determine Rh blood group (Liao et al., 2023). In both regions, the pangenome graph is too complex to be visualized in full; to simplify it, we pruned variant edges except for those with very long alleles (combined length >1kb), deleted ‘tips’ not contained in a cycle, and contracted non-branching paths (see Supplementary Note, Section 11). The graph was visualized as a bandage plot (Wick et al., 2015). This approach relies on our variant definition because there is no similar way to simplify the graph without first discarding unwanted edges.

The HLA-A region contains 4,704 total variants in just 3 superbubbles, which are visually apparent in the simplified graph (**Figure 4a**). The three superbubbles contain 75, 68, and 24 alleles, with average allele lengths of 53kb, 20kb, and 66kb, respectively; however, most of these alleles are highly similar, differing only by small variants. In the simplified graph, there are only three structurally different walks through the first two superbubbles, and two through the third. These correspond to large structural variants: the first bubble is a deletion of a 55kb region containing a nested 1kb deletion; the second is a near-deletion of a 21kb region also containing a nested 1kb deletion; the third is a 66kb insertion. The 66kb insertion, which contains the HLA-Y pseudogene, contains 98 variants and 24 unique alleles; all of these variants are non-GRCh38 and are therefore missed by *vcfwave*.

**Figure 4:**
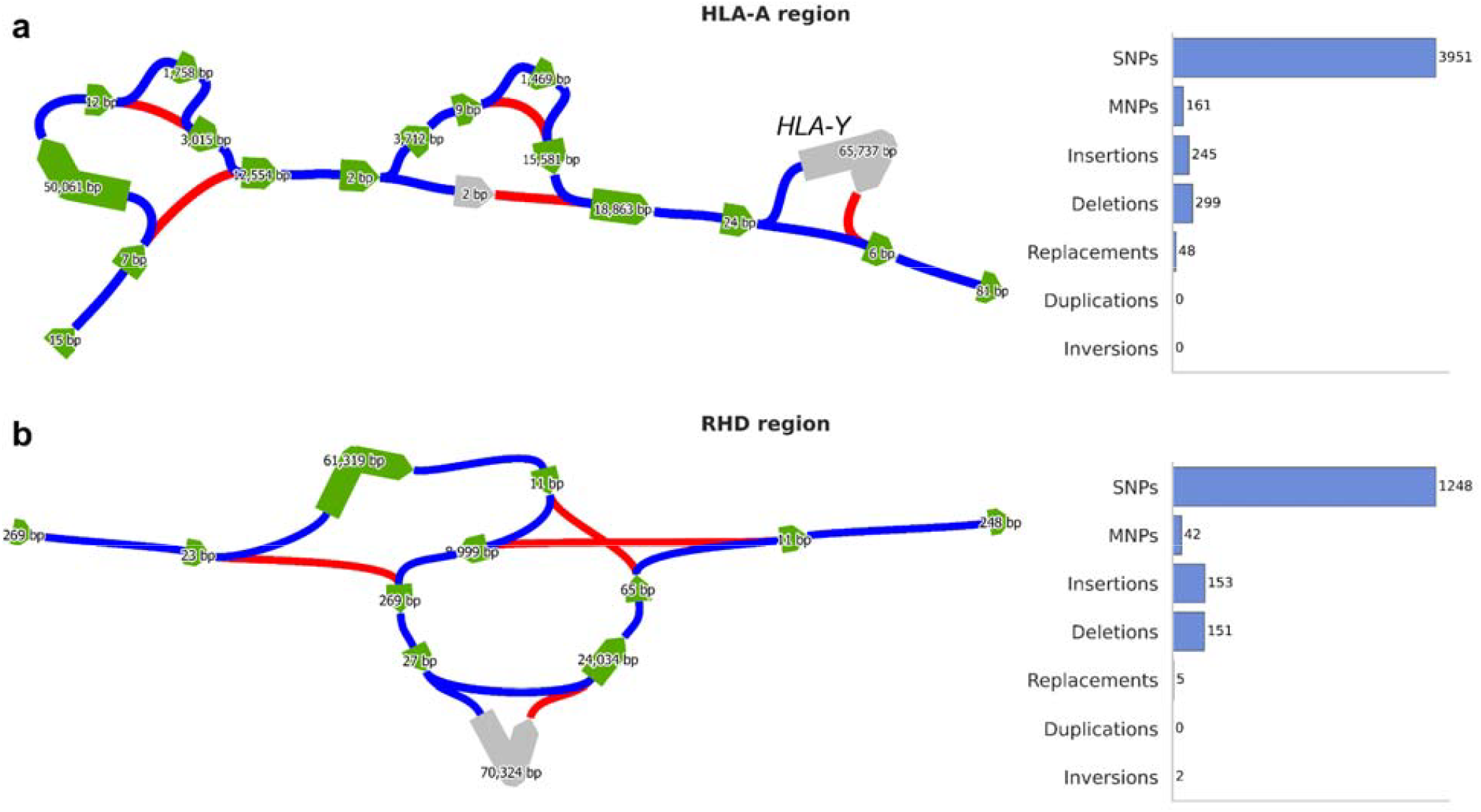
Analysis of two complex regions. Blue and red edges indicate reference tree paths and paths containing variant edges, respectively. Green and grey nodes indicate GRCh38 and non-GRCh38 sequences, respectively. (a) Simplified graph of the HLA-A region and a summary of variants identified. The location of the HLA-Y pseudogene is indicated. (b) Simplified graph of the RHD region and a summary of variants identified.

The RHD region forms a single superbubble containing 86 unique alleles (average length: 8.4kb); we identify 1601 variant edges, 78% of which are SNPs. Only one haplotype carries the inversion, which is 33kb long. With our definition, an inversion is represented by a pair of inversion edges, both of which are carried by that haplotype (see Discussion). The region also harbors a common 70kb deletion of the RHD gene (alternative allele count: 27) and a common 70kb insertion which duplicates the RHD gene (alternative allele count: 6). These variants differ in length by 5bp, and they are spaced 296bp apart. The insertion is nested inside of the inverted region; the deletion is interlocking with it, spanning one of the two inversion break points.

This kind of analysis can be done manually, but it is greatly streamlined by our approach, which makes it easy to enumerate structural variants and visualize their relationships. Fine-scaled variation can be removed for visualization and then mapped back to the simplified graph.

## Discussion

The human pangenome is intended to address the problem of reference bias, that no single reference genome is an adequate point of comparison for all other genomes. Indeed, the pangenome graph contains 129.3 megabases of non-GRCh38 genetic sequence (Liao et al., 2023); however, existing definitions of a genetic variant in the pangenome still rely heavily on the linear reference. They are unable to resolve differences - even SNPs - between different non-reference sequences (i.e., non-GRCh38 variants). We addressed this challenge by replacing the linear reference genome, which lacks non-GRCh38 sequences, with a reference tree containing all nodes and sequences in the graph. With this approach, we found 3.5 million genetic variants that are not located on the linear reference genome.

The linear reference genome, our reference tree, and the pangenome reference graph are all referred to as a “reference”. We suggest that the reference tree is logically parallel to the reference genome, whereas the pangenome itself is logically parallel to a reference *panel* like 1000 Genomes. A reference tree, like a reference genome, is a point of comparison; a pangenome, like a reference panel, contains both reference and alternative sequences. Any haplotype in a reference panel can be reconstructed by starting from the reference genome and replacing reference with alternative alleles; likewise, a haplotype in a pangenome graph can be reconstructed by starting from the reference tree (see Supplementary Note, Section 2).

Just as the choice of reference genome always affects the way that genetic variants are represented - including their REF vs. ALT alleles, and their positions - the choice of reference tree affects the variant edges that are found. The total number of variants is fixed, but not the variants themselves, and also not the number of SNPs or indels. What does remain constant is the difference between any two haplotypes and the fact that their difference is encoded uniquely as a combination of variant edges.

We published our variant catalogue in a VCF file in order to streamline downstream applications, but this choice required certain compromises or workarounds. The reference allele (REF) column in a VCF is required to match the reference genome at the specified position, so the reference allele of a non-GRCh38 variant must be written in a different field. Moreover, in a VCF, insertions and deletions have a letter prepended to both their reference and alternative alleles; we follow this convention for variants on the linear reference but not for variants off of it, to avoid breaking the reference allele rule. The position (POS) column of a VCF is a single integer, whereas we define variant position as a tuple, one of which must be chosen for the POS column. POS encodes the position of the first letter of each allele, which is offset by one from the way that we would choose to define it, which is the last letter before each allele begins (this approach more gracefully handles insertions). In each of these cases, we added custom INFO fields to the VCF file such that no information is lost, but users of our VCF files should be aware of these workarounds, especially when analyzing non-GRCh38 variants. For example, users should not misinterpret non-GRCh38 variants as being insertions just because their REF column is empty; we do include a variant type (VT) field (see Methods).

Inversions are particularly challenging to analyze in a pangenome graph. Defining inversions requires orienting the nodes of the bidirected graph (Minigraph-Cactus assigns arbitrary orientations). A choice of reference tree implies a choice of orientations; the same variant edge could be an inversion with respect to one reference tree and not with respect to another. Whereas most variant edges likely correspond to a single mutational event, an inversion event corresponds to a pair of variant edges with opposing orientations: from the positive to the negative orientation, and back. Sometimes inversion edges can be paired up in this way, but not always, in particular when the inverted region overlaps with an assembly gap. Therefore, we represent inversion edges individually, without assuming perfect pairing.

Although we suggest that superbubbles should no longer be used to define genetic variants, they are useful to highlight complex regions. Simply counting either variant edges or alleles per superbubble is a useful heuristic for the complexity of a genomic region, with complex regions often having many alleles and variants. Indeed, a large fraction of non-GRCh38 variants are contained in highly multiallelic superbubbles. When a variant is nominated for special scrutiny, for any reason, a reasonable starting point would be to analyze its enclosing superbubble; one could enumerate variant edges in that superbubble, visualize it (perhaps after simplification), or even inspect its alleles by eye.

Whereas the GRCh38 reference genome forms a simple path - not visiting any node twice - in the Minigraph-Cactus graph, it does not do so in the pangenome graph inferred using PGGB, and other reference genomes (e.g., CHM13) do not generally form a simple path in either graph. Our framework is capable of handling linear reference genomes with repeated nodes (see Supplementary Note, Section 12), but not without complications. In particular, nodes that belong to a reference-genome cycle no longer have a single position, and multiple reference positions are superimposed in the graph. We prefer to define variant positions against an acyclic reference genome path because if a reference sequence is duplicated, and subsequently a mutation arises on one copy, then it is most natural to define the position of that mutation in relation to that copy specifically.

## Methods

### Analysis Details

#### Chromosome-specific graph extraction

We extracted subgraphs for chromosomes 1–22, X, and Y from the full HPRC .gfa file (see Data Availability). For each chromosome, we identified the node corresponding to its first occurrence in the graph and used gfatools to extract the chromosome-specific subgraph. This per-chromosome segmentation enabled parallelization and reduced memory requirements.

### Exclusion of terminal variant edges

To construct the reference tree, we introduced four terminus nodes (two binodes) to anchor disconnected regions (see Supplementary Note, Section 3). Variant edges involving these termini are called terminal edges, and they are a convenient mechanism to track the start and end of each walk. Terminal edges mostly do not represent biological variation, but rather the presence of alignment gaps, and they were excluded from both our published VCFs and our variant count statistics. More precisely, for a variant edge (*u, v*), if either the branch point of *u* and *v* or the node *v* is one of the terminus nodes, then (*u, v*) is discarded.

#### Reference and alternative allele count

Each sample in the GFA file is represented as a collection of walks. The reference allele count for a variant is defined as the number of times a walk visits the last edge of corresponding reference path (see Supplementary Note, Section 7) in the reference tree. The alternative allele count is defined as the number of times a walk traverses the variant edge.

#### Non-GRCh38 variants

The variant edge (*u, v*) was classified as a non-GRCh38 variant if no GRCh38 walk visits *v*.

#### Repeat Annotation

To annotate local tandem repeats and homopolymeric motifs, we identified indels such that (1) their non-empty allele is (*s*)^*n*^ for some sequence motif *s*, and either (2a) the upstream sequence in the reference tree equals *s*, or (2b) some downstream sequence in the reference tree equals s. This approach recovers tandem repeat variants in cases where the minigraph-cactus graph topology lacks repeat structure (see Supplementary Note, Section 9). It can potentially miss copy number changes of *s* ➔ (*s*)^2^ or (*s*)^2^ ➔ *s* when the matching motif is split between upstream and downstream sequences.

#### Region Annotation

We used BED files from (Dwarshuis et al., 2023). Each variant was labeled as belonging to one of three categories: Easy, Segmental Duplication (Segdup), or Hard regions. Regions not explicitly labeled as Easy or Segdup were designated Hard. We annotated variants based on their VCF positions, excluding variants in segmental duplications from the hard category, such that the three categories formed a partition.

#### Variant Length Stratification

Variants were categorized as either large or small using their total allele length, defined as the sum of the reference and alternate allele lengths, and a threshold of at least 50 basepairs.

#### vcfwave

*vcfwave* is an approach that analyzes multiallelic superbubbles in a pangenome graph. For each superbubble, it first identifies all possible alleles - typically long sequences corresponding to distinct haplotype walks through the superbubble. These alleles are then realigned to one another to detect finer-scale variation within the superbubble, such as SNPs, small indels, or nested variant structures. This realignment-based strategy enables *vcfwave* to decompose complex, long variants into smaller, interpretable events within complex regions (Liao et al., 2023).

#### Insertion and Deletion Superbubbles

A superbubble is defined as an insertion if the first node and the last node of the superbubble are adjacent via a reference-tree edge. Similarly, a superbubble is defined as a deletion if the first node and the last node of the superbubble are adjacent via a variant edge.

#### Degenerate Alleles

To maintain clarity in variant representation, we excluded exactly degenerate alleles (i.e., variants with identical REF and ALT alleles) from the VCF and downstream analyses.

#### Performance and Resource Usage

Our pipeline was parallelized by chromosome. Chromosome 1 required 36 GB of memory and 10 hours of wall time.

### VCF Specification

Our variant call format (VCF) extends the standard specification, maintaining compatibility with conventional tools while also including pangenome-specific information.

#### Variant Identifiers and Canonical Fields

Each variant is identified directly by its corresponding edge in the pangenome graph, replacing the traditional bubble ID convention. The POS, REF, and ALT are preserved, but with critical reinterpretations:

- CHROM: The chromosome number.
- POS: The VCF position, which is a single integer, differs from our variant-edge position, which is a pair of integers (see ‘PU’ and ‘PV’ below). For indels such that we prepend a letter to both alleles, the VCF position of the variant edge (u,v) is the position of node u. Otherwise, it is one plus the position of u. The VCF is sorted based on POS.
- ID: The variant edge ID is a string concatenating the edge’s node IDs of the pangenome graph from the original GFA file. Each node ID contains either ‘>’ or ‘<’ symbols that indicate the orientation of the node in the Pangenome graph (‘>’ means positive orientation and ‘<’ means negative orientation). It should be noted that these orientations are inherited from the pangenomes GFA file, therefore, they might be different from the orientation of the reference tree as we orient the nodes (see Supplementary Note, Section 5).
- REF/ALT: Each variant edge corresponds to a pair of alleles: the reference allele derived from the reference tree, and the alternate allele from the non-reference path. The alternate allele is reported in the ALT field. If the reference allele lies entirely on the GRCh38 linear reference path, it is recorded in the standard REF field in the VCF. If the reference path does not align with the GRCh38 reference, the REF field is set to ‘.’ because REF is required to match the linear reference genome. Non-GRCh38 reference alleles are recorded in the NR (non-reference reference allele) field (see below).

For indels on the linear reference, we follow the VCF convention that one letter - the letter preceding the indel on the linear reference - is prepended to both REF and ALT. We do not do so for non-GRCh38 indels, whose reference allele is recorded in the NR field instead of REF.

- QUAL: Arbitrarily set to 60.
- FILTER: Set to ‘PASS’.
- INFO (**Customized VCF Fields)**. The INFO field includes the following items:
  - **NR** (Non-Reference allele): The allele obtained by traversing the reference tree from the branch point to the end of the variant edge. If this path is off the GRCh38 linear reference, REF is set to ‘.’ and NR holds the reference-tree-based allele (see Supplementary Note, Section 7). Otherwise, NR is ‘.’.
  - **VT** (Variant Type): Annotates the variant as one of the following: SNP, MNP, insertion, deletion, replacement, duplication, or inversion.
  - **DR** (Distance from Reference): A tuple capturing the distance of each node in the variant edge from the linear reference path (GRCh38).
  - **RC** (Reference allele Count): Total number of times all haplotype walks traverse the corresponding reference tree edge.
  - **AC** (Alternative allele Count): Total number of times all haplotype walks traverse the variant edge.
  - **AN:** Total number of times all haplotype walks traverse either the variant edge or the corresponding reference tree edge.
  - **PV** (Positions of variant nodes): A tuple storing the positions of nodes u and v in the variant edge (u, v).
  - **TR_MOTIF:** Contains the repeated motif if the variant edge corresponds to a local tandem repeat, otherwise ‘.’.
  - **NIA (Nearly Identical Alleles):** Nearly identical alleles are defined as allele pairs ≥10 bp differing by only one base (through substitution or a 1-bp indel), were flagged but retained. NIA is either ‘yes’ or ‘no’ flags nearly identical alleles.
- **Genotype data**. Each individual’s phased genotype is represented (sample ID) using a structured format in the sample-level fields: GT : CR : CA.
  - **CR / CA**: Respectively count the number of times a haplotype visits the reference tree edge and the variant edge. The reference edge is defined as the last edge of the reference path. Because haplotype assemblies consist of multiple contigs represented as disconnected walks, some variant edges may not be visited simply due to gaps in assembly. In such cases, if the position interval of a variant edge falls entirely outside the haplotype’s linear coverage, both CR and CA are marked as missing (.) in the VCF (see Supplementary Note, Section 10).
  - **GT** (Genotype): Indicates the presence or absence of the variant edge and the reference edge. GT is 1 if CA is positive, 0 if CA is 0 and CR is positive, and ‘.’ if both are missing.
  - **Missingness:** A variant is considered missing in a haplotype if all three fields (GT, CR, and CA) are ‘.’. This position-based strategy efficiently approximates true missingness without requiring exhaustive graph traversal (see Supplementary Note, Section 10).

### Comparison with Bubbles

#### Assignment of Variant Edges to Superbubbles

We used published .snarls files (see Data Availability) to assign variant edges to superbubbles. This file includes bubbles of levels 0 to 28 and defines each bubble by listing its constituent nodes and an associated ID. Superbubbles are level 0 bubbles. A variant edge was assigned to a superbubble if both of its nodes were present within the same superbubble; we confirmed that every variant edge is assigned to exactly one superbubble. This approach was applied genome-wide across all 22 autosomes, as well as chromosomes X and Y. We found that every variant edge was contained within a superbubble, except those involving artificial terminus nodes used for reference tree construction.

We did not compare against the full hierarchical bubble structure (levels ≥1), for two primary reasons. First, higher-level bubbles do not yield variant definitions with well-defined positions along the linear reference genome, limiting interpretability and compatibility with VCF-based representations. Second, these bubbles introduce redundancy: the same variant may appear in multiple bubbles across different levels, each offering a different representation, which complicates downstream analysis.

#### SNP Benchmarking Against External VCFs

We compared SNPs identified in our output VCF with those in the original HPRC VCF and vcfwave. SNPs were matched based on position, reference allele, and alternative allele. Variants found in both datasets were considered shared, while those present in only one were labeled as dataset-specific. This comparison was limited to SNPs because for non-SNPs, matching can be ambiguous; for example, a deletion in a homopolymer can be coded in several different ways, with different positions but equivalent sequence effects.

#### Classification of Triallelic Structures

We defined a bubble as triallelic if it contained two variant edges. Triallelic bubbles were categorized into four types: properly triallelic, overlapping, nested, and interlocking. The classification was based on the two branch points of the variant edges within the bubble and the two end nodes of the bubble:

Properly triallelic: Bubbles where both start and end nodes have degree three.

Overlapping: Bubbles where exactly one of the start or end node has degree three.

Nested: Bubbles where one of the variant edges has the same branch point as the starting node of the bubble and end node as the last node of the superbubble.

Interlocking: Bubbles where one of the variant edges has the same branch point as the starting node of the bubble and the other variant edge has the same end node as the last node of the superbubble.

### Complex Regions

#### Region Preprocessing and Positioning

We extracted subgraphs corresponding to specified genomic intervals of interest. These intervals were selected based on their known structural complexity or biological relevance, and their positions are reported in **Supplementary Table S11**.

The genomic coordinates for the HLA and RHD regions were obtained from the NCBI Genome database (hg38). Using these coordinates, we manually identified potential superbubbles corresponding to the target genes by locating the relevant node IDs in the generated VCF file. We then visualized these regions using Bandage and compared the resulting graph structures to those presented in the first draft of the human pangenome reference (Liao et al., 2023) to refine the exact node IDs and positions associated with the HLA and RHD loci.

#### Simplification of Complex Variant Structures

To enable effective interpretation of complex genomic regions, particularly those containing bubbles with dense clusters of small variants, we applied a graph simplification procedure. Variant edges with short combined allele lengths were filtered out, tip-like structures not contributing to connectivity were pruned, and linear paths comprising degree-two nodes were contracted (see Supplementary Note, section 11). The resulting graph retained long, structurally informative variants. They were visualized as bandage plots using an existing tool (see Code Availability).

## Supporting information

Supplementary Tables List

## Data availability

We have released genome-wide variant call format (VCF) files generated using our variant edge framework. These include VCFs based on both the GRCh38-aligned Minigraph-Cactus V1.1 pangenome graph and the CHM13-aligned Minigraph-Cactus graph. All VCFs are publicly available at Zenodo: https://zenodo.org/records/15374896. The files include full-genome VCFs for all autosomes and chromosomes X and Y with per-individual genotypes.

The pangenome graphs were constructed by the Human Pangenome Reference Consortium (HPRC) using Minigraph-Cactus v1.1, and are available as .gfa files at: https://s3-us-west-2.amazonaws.com/human-pangenomics/pangenomes/freeze/freeze1/minigraph-cactus/hprc-v1.1-mc-grch38/hprc-v1.1-mc-grch38.gfa.gz. Superbubble annotations were obtained using .snarls files precomputed by the VG toolkit for the Minigraph-Cactus v1.1 graph. To stratify genomic regions by complexity, we used BED files defining “easy,” “segmental duplication (segdup),” and “hard” regions downloaded from: https://ftp-trace.ncbi.nlm.nih.gov/giab/ftp/release/genome-stratifications/v3.5/GRCh38@all/Union/GRCh38_notinalldifficultregions.bed.gz (easy region), https://hgdownload.soe.ucsc.edu/goldenPath/hg38/database/genomicSuperDups.txt.gz (segdup region).

## Code availability

We have released an open-source software package, pantree, which implements our reference tree construction, variant edge definition, and VCF generation pipeline. The codebase is implemented in Python and is available at https://github.com/oclb/pantree. Scripts are available at https://github.com/ShenghanZhang1123/graph_var_analysis.

In addition to our own code, we used several publicly available tools in this study. The VG toolkit (https://github.com/vgteam/vg) was used to extract superbubbles information from .snarls files and detect superbubbles within the pangenome graph. We used gfatools (https://github.com/lh3/gfatools) for general graph manipulation and format conversion tasks. For visualization of pangenome graph structures, we used Bandage (https://github.com/rrwick/Bandage). All tools are open-source and freely accessible.

## Acknowledgements

L.J.O acknowledges funding under NIH-NIGMS award R35-GM155278 and Simons Foundation SFARI award 1009802LO. H. L acknowledges funding under NIH-NHGRI award R01-HG010040, U01-HG013748, and U41-HG010972.

## Author contributions

P.S.N., S.Z., H.H., H.L., and L.J.O developed the methods. S.Z. and H.H. performed the experiments. P.S.N., S.Z., H.H., H.L., and L.J.O. wrote the paper. H.L. and L.J.O. supervised the research.

