## Supplementary Note for "Defining and cataloging variants in pangenome graphs"

This supplementary note presents the mathematical and algorithmic foundations of our framework for defining genetic variants in pangenome graphs. The input is a bidirected graph constructed from a collection of walks, each representing a sample haplotype. Bidirected graphs naturally capture the double-stranded nature of DNA, allowing traversal from either orientation at each node. The central mathematical question is: given such a graph, can we define a minimal set of features that uniquely encodes all pairwise differences between walks? We formalize this in terms of symmetric differences and show that a set of variant edges, non-tree edges with respect to a reference tree, serves as a basis for all such differences. This document details our definitions, algorithms, and theoretical guarantees, including walk reconstruction, variant positioning, and variant edge classifications.

#### 1 Pangenomes as Bidirected Graphs

Pangenome graphs are naturally modeled using bidirected graphs due to the strandedness of DNA, which requires representing both directions of sequence traversal. Throughout this work, we define a *bidirected graph* $B = (V, E, C)$ via its *directed representation*:

- $V$ is the set of nodes,
- $E \subseteq V \times V$ is the set of directed edges,
- $C : V \to V$ is a fixed-point-free involution called the complement function, satisfying $C(v) = v^c$, $(v^c)^c = v$, and $v \neq v^c$ for all $v \in V$,
- For every edge $e = (u, v) \in E$, its complement is defined as $e^c := (v^c, u^c) \in E$, so every edge appears in a pair with its directed complement.

Using these criteria, a bidirected graph can be represented as a directed graph.
The *bidirected representation* of $B$ is the quotient structure $(V/C, E/C)$:

- A *binode* is an unordered pair $\{u, u^c\} \in V/C$,

- A *biedge* is an unordered pair $\{e, e^c\} \in E/C$.

This abstraction collapses each node with its complement and each edge with its reverse complement. For visualization purposes, we depict a binode as a double-headed arrow, where each side represents a direction of traversal. Figure 1.A illustrates the bidirected representation: each binode is shown as a double-headed arrow, and biedges connect the heads or tails depending on the direction of entry and exit. In this figure, traversing from the left side to the right side of a binode corresponds to reading the forward strand sequence, while traversing from right to left corresponds to the reverse complement. Figures 1.B and 1.C show the directed representation of the same bidirected graph, where each binode is "unfolded" into a pair of nodes $v^+$ and $v^-$, and each biedge becomes a pair of directed edges. In this representation, forward and reverse traversals correspond to complementary nodes.

These two representations are mathematically equivalent and are used interchangeably depending on context.

A *walk* between binodes $\{v_1, v_1^c\}$ and $\{v_n, v_n^c\}$ is defined as a pair of node sequences:

$$(v_1, \ldots, v_n) \quad \text{and} \quad (v_n^c, \ldots, v_1^c),$$

such that each pair $(v_i, v_{i+1}) \in E$ for $1 \leq i < n$. A walk is called a *path* if it does not repeat any binode: that is, $\{v_i, v_i^c\} = \{v_j, v_j^c\}$ implies $i = j$.

A bidirected graph is a *rooted tree* with root $\{r_1, r_1^c\}$ if for either $r_1$ or $r_1^c$, say $r$, and for all binodes $\{v_1, v_1^c\}$, there exists a unique directed path from $r$ to either $v_1$ or $v_1^c$. Such a bidirected graph is *oriented*: its nodes $V$ can be partitioned into two components with no edges between them, $V^+$ and $V^-$, such that if $v \in V^+$ then $v^c \in V^-$ (see Section 5).

A bidirected graph $S = (V_s, E_s, C)$ is a *subgraph* of a bidirected graph $B = (V, E, C)$ if $S$ is a bidirected graph and $V_s \subseteq V$ and $E_s \subseteq E$ i.e., $V_s$ and $E_s$ are closed under complement i.e., if $v \in V_s$ then $v^c \in V_s$ and if $e \in E_s$ then $e^c \in E_s$. A subgraph $S$ is called a *spanning subgraph* if $V_s = V$. A *rooted spanning tree* of a bidirected graph is a spanning subgraph that is a rooted tree.

### 2 Reference Tree and Variant Edges

Given the pangenome graph $B$ and a rooted spanning tree $T$, called a *reference tree*, we define *variant edges* as the biedges of $B$ that are not in $T$. Let $T$ be a reference tree of $B$, then $T$ induces two directed trees in the directed representation of $B$ called *half-trees* such that one of them has edges only from $V^+$ to $V^+$, denoted by $T^+$, and the other one only has edges from $V^-$ to $V^-$, denoted by $T^-$.

The representative of a variant biedge $\{e, e^c\}$ is either $e$ or $e^c$. For consistency, if $\{e, e^c\} = \{(u^+, v^+), (u^-, v^-)\}$ we pick $(u^+, v^+)$ as the representative. If a variant edge is between $V^+$ and $V^-$, it is an *inversion* edge. Note that if an inversion edge $e$ goes from $V^+$ to $V^-$, then $e^c$ also goes from $V^+$ to $V^-$, and similarly for an inversion edge from $V^-$ to $V^+$, its complement also goes from $V^-$ to $V^+$. For inversion edges, we randomly choose the representative.

The *genotype* of an individual, represented as a collection of disjoint walks through the graph, is the number of times that the walks visits each variant edge. We use the notation $x_e(W)$ for the number of times that walk $W$ visits biedge $e$.

Given the sequence of variant edges visited by walk $W$, as well as its starting and ending nodes, $W$ can be reconstructed uniquely because there is at most one path between any two nodes in the reference tree, in particular from the ending point of one variant edge to the starting point of the next one. Moreover, given the genotype of a walk – which tracks variant edge visit counts, but not their order – it is possible to reconstruct the number of times that it visits every biedge.

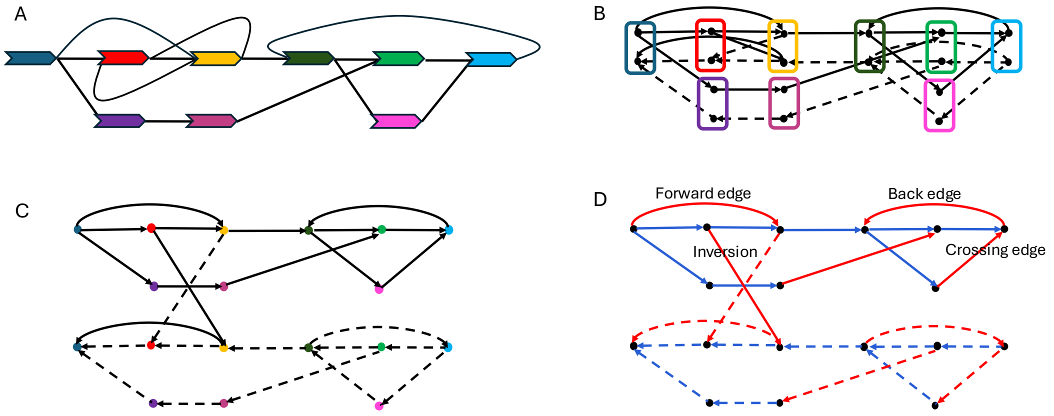


Figure 1: **Bidirected and directed representations of a pangenome graph.** (A) Bidirected representation of a bidirected graph. Each binode is shown as a double-headed arrow. Biedges connect either heads or tails depending on traversal orientation. For example, the biedge between the purple and dark pink binodes connects the forward orientation of the purple binode to the forward orientation of the dark pink binode, and the reverse orientation of the dark pink binode to the reverse orientation of the purple binode. (B) Directed representation of the same graph, where each binode is unfolded into two complementary nodes, one representing forward orientation and one representing reverse orientation. The color scheme follows panel (A). Each biedge corresponds to a pair of directed edges—one shown solid, one dashed. (C) An expanded version of (B), where complementary nodes are spatially separated for clarity. (D) The reference tree highlighted in blue. The tree over forward oriented nodes forms a DFS tree. Variant edges (in red) are classified relative to this tree as forward, back, or crossing edges. Inversion edges connect a forward oriented node to a backward oriented node.

**Theorem 2.1.** *Given a bidirected pangenome graph $B$, its reference tree $T$, and the starting and ending binodes of a walk $W$, the genotype $x(W) = \{x_e(W) \mid \text{variant edges } e\}$ determines $W$ up to the order of cycles.*

Before proving Theorem 2.1, we state and prove the following lemma.

**Lemma 2.2.** *Let $B$ be a bidirected graph with a reference tree $T$, which induces the partition $V = V^+ \cup V^-$. Let $W, W^c$ be a bidirected walk in $B$. Let $v_0, u_0$ be the starting and ending nodes of $W$, respectively, and let the variant edges visited by $W$ be $(u_1, v_1), \ldots, (u_n, v_n)$. For $i = 0, \ldots, n$, call $v_i$ a source if $v_i \in V^+$, and otherwise call $v_i^c$ a sink; if $u_i \in V^+$ call it a sink, otherwise call $u_i^c$ a source. Let sources$(W)$ and sinks$(W)$ denote the sequences of sinks and sources. For a node $v \in V^+$, let $D(v) \subset V^+$ be its reach set in $T$. Then the number of times that $W$ visits the binode $\{v, v^c\}$ is:*

$$|D(v) \cap \mathit{sinks}(W)| - |D(v) \cap \mathit{sources}(W)|.$$

*Proof.* We decompose $W$ into a sequence $S_0, ..., S_n$ of directed paths on $T$, where $S_i$ proceeds from the $i$th source node to the $i$th sink node. Each path in this sequence starts at a source node, ends at a sink node, and remains within $V^+$. Each path is a subsequence either of $W$ or of $W^c$.

If $S_i$ does not visit $v$ then either:

- It starts and ends outside $D(v)$ (non-descendant to non-descendant), or
- It starts and ends within $D(v)$ (descendant to descendant),

but it cannot go from a descendant to a non-descendant, and if it goes from a non-descendant from a descendant then it would visit $v$.

Therefore, the number of paths in $S$ that visit $v$ equals the number of transitions into $D(v)$:

$$|D(v) \cap \text{sinks}(W)| - |D(v) \cap \text{sources}(W)|.$$

Moreover, the number of paths that visit $v$ is equal to the number of times that $W$ (or $W^c$) visits either $v$ or $v^c$. □

*Proof of Theorem 2.1.* The visit count formula in Lemma 2.2 is invariant when the order of sources and sinks is shuffled; any two walks with the same multiplicity of sources and sinks visit a binode $b$ the same number of times. The multiplicity of sources and sinks can be computed from the genotype of a walk, so the genotype determines the visit count of $b$. It also determines the number of times each tree edge is traversed. The only ambiguity is within cycles, i.e., in the relative order of subpaths that begin and end at the same binode.

Thus, the genotype $x(W)$ determines the walk $W$ up to the order of cycles. □

To demonstrate the practical application of Theorem 2.1, we now present Algorithm 2.3, which efficiently constructs the bidirected walk $W$ from the genotype $x(W)$.

**Algorithm 2.3.** *The algorithm reconstructs the walk using the endpoints of the walk and the genotype information:*

1. ***Determine Sources and Sinks:*** *Sources and sinks are determined based on the variant edges. For each visit to a variant edge $(u, v)$, if $u$ is a positive node, it is added to the sinks; if it is negative, its complement is added to the sources. Similarly, if $v$ is positive, it is added to the sources; if it is negative, its complement is added to the sinks.*

2. ***Adjust for Inversions:*** *The start and end nodes of the walk are added to the sources or sinks depending on the number of inversions. If the number of inversions is even, the start node is added to the sources and the end node to the sinks. If the number of inversions is odd, additional adjustments are made: if there is one more* $+$ *to* $-$ *switch, both endpoints are added to the sinks; if there is one more* $-$ *to* $+$ *switch, both endpoints are added to the sources. After doing this, there are an equal number of sources and sinks (otherwise, the genotype does not correspond to a valid walk).*

3. ***Reconstruction of the Walk:*** *The reconstruction proceeds by traversing the positive-orientation reference tree in the opposite direction of the tree edges. Starting from each sink, the algorithm walks upward along the tree until reaching a source, recording each edge traversed along the way. This process avoids searching multiple branches as every node has at most one predecessor, but any number of descendants. The number of times each edge is visited is stored. When a source is reached, its count is decremented (a source is used up when its count reaches* $0$*). This process is repeated for all sinks, ensuring all sources are matched, thus reconstructing the walk up to the order of cycles.*

4. ***Validation of the Genotype:*** *This algorithm also detects whether the genotype is valid: if it ever reaches the root, and the root no longer belongs to the set of sources, then the genotype corresponds to no single walk.*

### 3 Constructing the Reference Tree

Suppose the bidirected graph $B(V, E, C)$ is given. Before finding the reference tree, we prespecify the direction of each node in positive and negative classes, denoted $V^+$ and $V^-$ respectively (see section 5) meaning that if f $v \in V^+$, then $v^c \in V^-$, and vice versa. To ensure that $B$ has a rooted spanning tree, we add two special binodes to $B$, called the start-terminus and the end-terminus, denoted by ST and ET. We also add the biedge $((\mathrm{ST}^+, v^+), (v^-, \mathrm{ST}^-))$ if $v^+ \in V^+$ either has in-degree zero in $B$ or is the starting node of a walk. Recall that $B$ is constructed from a collection of walks, where each sample haplotype consists of a collection of walks. Similarly, for a binode $(v^+, v^-)$, if $v^- \in V^-$ either has in-degree zero or is the end of a walk, we add the biedge $((\mathrm{ET}^-, v^-), (v^+, \mathrm{ET}^+))$. After adding these nodes, there exists a directed spanning tree of the positive subgraph with root $\mathrm{ST}^+$. This tree along with its complement on the negative subgraph forms a spanning rooted tree for the bidirected graph $B$. To construct this graph, we use a DFS traversal on the positive subgraph initiated at $\mathrm{ST}^+$. This tree and its complement are our reference tree.

If the pangenome graph includes the linear reference as a path, with no repeated binodes, then we can ensure that nodes connected via the linear reference have the same direction. We can initialize our DFS traversal at this path. This is the case for the minigraph-cactus graph, but not the PGGB graph. It is desirable to include the linear reference as the primary branch of the reference tree because it allows us to define genomic position (see section 4).

There exist many possible choices of reference tree, none of which is the best; Theorem 2.1 holds for any choice of reference tree. We use a heuristic designed to greedily maximize the allele frequency of reference alleles. When the DFS search has multiple options for its next step, it always chooses the edge with the highest frequency (i.e., the largest number of walks that visit that edge).

#### 4 The Position of Variants Along the Linear Reference

In practice, it is important that the variants have a well-defined position with respect to the linear reference genome. We assign positions to variants by making an appropriate choice of reference tree. In particular, we choose a depth-first search (DFS) tree whose first branch is the linear reference genome. If a node $u$ is not on the linear reference, it is assigned the largest position of any node on the linear reference that can reach $u$ in $T$. Equivalently, this is the largest position of any node on the linear reference that can reach $u$ in $B$ without visiting a back-edge with respect to $T$.

The position of a binode $\{u, u^c\}$ is defined as the linear reference position of $u$ (or $u^c$) if either one lies on $L$. Otherwise, it is the position of the node $x$ on $L$ that serves as the ancestor of $u$ (or descendant of $u^c$) in $T$. The position of a variant edge $e = (u, v)$ in $B$, denoted by $\text{pos}(e)$, is defined as the interval $[a, b]$, where $a$ is the position of $u$ and $b$ is the position of $v$.

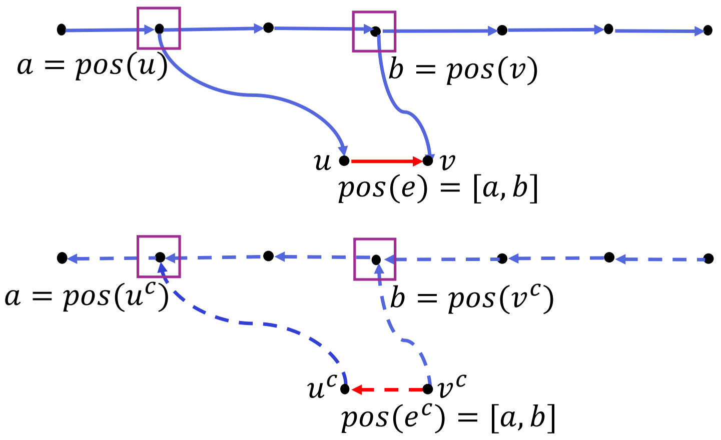


Figure 2: **Position of a variant edge and its complement.** The blue paths represent the linear reference in forward (solid) and reverse (dashed) orientations. The position of a node $v$ is defined as the position of its lowest ancestor in the forward-oriented linear reference (or its first descendant in the backward-oriented reference). For a variant edge $e = (u, v)$, the position is defined as the interval $\text{pos}(e) = [\text{pos}(u), \text{pos}(v)]$. The position of a node is the same as that of its complement, and similarly, the position of a variant edge is equal to the position of its complementary edge.

**Theorem 4.1.** *For an inversion-free bidirected graph, if the position of $e$ is $[a, b]$, then $x_e(w)$ is determined by the variant edges whose position overlaps $[a, b]$:*

$$\{x_\eta(w) \mid \textit{variant edges } \eta \textit{ s.t. } pos(\eta) \cap [a, b] \neq \emptyset\}$$

*Proof.* By the definition of position, each node not on the linear reference inherits its position from the highest ancestor on the linear reference reachable via $T$. Since there is a path on $T$ from $u$ to $v$, by the definition of position $pos(u) \leq pos(v)$.

Therefore, if a path from $a$ to $b$ exists in $T$ and $\text{pos}(a) \neq \text{pos}(b)$, then the path must pass through the ancestor of $b$ that lies on the linear reference and has position $\text{pos}(b)$. □

*Proof.* Suppose $B$ is an inversion-free bidirected graph with a reference tree $T$. Consider a variant edge $(u, v)$ and a binode $b$ in $B$. Utilizing Lemma 2.2, we show that if $\text{pos}(v) \notin [\text{pos}(u), \text{pos}(v)]$, then for any walk $W$,

$$|D(b) \cap \text{sinks}(w)| - |D(b) \cap \text{sources}(w)|$$

remains invariant.

Without loss of generality, we may assume that $\text{pos}(u) \leq \text{pos}(v)$ since we only work within $T$ for the remainder of the proof. There are two cases:

**Case 1:** $\text{pos}(b) < \text{pos}(u)$.

- **Subcase 1.1:** $u \in D(b)$. In this case, there is a path within $T$ from $b$ to $u$. This path must pass through the ancestor of $u$ within the linear reference whose position is equal to $\text{pos}(u)$. Since $\text{pos}(u) \leq \text{pos}(v)$ and $(u, v)$ is not an inversion, there is a path from $b$ to the ancestor of $v$ within the linear reference whose position is equal to $\text{pos}(v)$, and therefore there is a path within $T$ from $b$ to $v$. Hence, $v \in D(b)$ and $|D(b) \cap \text{sinks}(w)| - |D(b) \cap \text{sources}(w)|$ remains unchanged.

- **Subcase 1.2:** $u \notin D(b)$. In this case, there is no path within $T$ from $b$ to the ancestor of $u$ within the linear reference whose position is equal to $\text{pos}(u)$. Therefore, since $\text{pos}(u) \leq \text{pos}(v)$ and $(u, v)$ is not an inversion, there is no path from $b$ to the ancestor of $v$ within the linear reference whose position is equal to $\text{pos}(v)$, and therefore there is no path within $T$ from $b$ to $v$. Hence, $v \notin D(b)$ and $|D(b) \cap \text{sinks}(w)| - |D(b) \cap \text{sources}(w)|$ remains unchanged.

**Case 2:** $\text{pos}(v) \leq \text{pos}(b)$. In this case, both $u$ and $v$ are not in $D(b)$, and therefore $|D(b) \cap \text{sinks}(w)| - |D(b) \cap \text{sources}(w)|$ remains unchanged. □

Theorem 4.1 shows that our definition of position not only has the linear nature of the genome, but also allows us to work locally: in order to reconstruct any walk through a locus, it suffices to consider those variants whose positions intersect with that locus.

The inversion-free assumption in Theorem 4.1 is essential. The following counterexample illustrates this point: Let $e = (u^+, v^+)$ be an edge with position $[a, b]$. Consider two inversion edges, $e_1 = (u_1^-, v_1^+)$ and $e_2 = (u_2^+, v_2^-)$, with positions $[a_1, b_1]$ and $[a_2, b_2]$, respectively. Suppose that $b_1 < a$ and $b < a_2$, so the positions of $e_1$ and $e_2$ do not intersect the position interval of $e$. Let $W_1$ be a walk that visits $e$ once, without visiting $e_1$ or $e_2$. Now let $W_2$ be a walk with the same genotype as $W_1$ except it visits both $e_1$ and $e_2$ exactly once. In this case, $W_2$ visits $e$ twice, despite having the same variant edge counts as $W_1$ within the interval $[a, b]$.

### 5 Orienting Nodes Within Each Binode

The reference is constructed via depth-first search on the positive-direction subgraph of the bidirected pangenome graph. Before doing so, we assign directions to each node of the graph.

A subgraph $P$ of a bidirected graph $B$ is a partition subgraph if it is a spanning subgraph and there is no connected component of the underlying undirected graph of $P$ containing both $v$ and $v^c$ for any $v \in V$. Such a graph induces a non-unique partition of $V$ into two components, labeled $V^+$ and $V^-$, with no directed edges between them, such that if $v \in V^+$, then $v^c \in V^-$.

A partition subgraph $P$ of a bidirected graph $B$ is called maximally connected if it has the fewest connected components out of all partition subgraphs.

The *undirected underlying graph* of a bidirected graph $B(V, E, C)$, denoted by $U(B)$, is defined as the undirected multigraph $U$ with vertex set $V(U) = V/C$ (i.e., $\{u, u^c\}$ becomes a single node), and edge set $(u, v) \in E(U)$ if either $u$ or $u^c$ is adjacent to either $v$ or $v^c$ in $B$.

**Proposition 5.1.** *For any bidirected graph $B$, a spanning forest of $U(B)$ that contains the minimum number of trees induces a maximally connected partition subgraph of $B$.*

*Proof.* Let $F$ be a spanning forest of $U(B)$ with the minimum number of trees. Define $\mathcal{F}$ to be the subgraph of $B$ whose edges correspond to those in $F$ (i.e., $\mathcal{F}$ is the restriction of $B$ to complementary biedges which correspond to the edge set of $F$). Then $\mathcal{F}$ is a spanning subgraph of $B$ whose connected components correspond to the trees of $F$.

Suppose a connected component $\mathcal{C}$ of $\mathcal{F}$ contains an edge $v$ with directed paths to (or from) both $u$ and $u^c$. Let $\mathcal{P}_1$ and $\mathcal{P}_2$ be these directed paths. By the definition of the underlying undirected graph, $C = U(\mathcal{C})$ is connected, and $U(\mathcal{P}_1) \cup U(\mathcal{P}_2)$ forms a cycle in $C$, which is a contradiction since $\mathcal{C}$ should not contain cycles. Therefore, $\mathcal{F}$ must be a partition subgraph of $B$.

Now, suppose there exists another partition subgraph of $B$, denoted $\mathcal{F}'$, with fewer connected components than $\mathcal{F}$. Then, $U(\mathcal{F}')$ would have fewer connected components than the number of trees in $F$, contradicting the assumption that $F$ is a spanning forest of $U(B)$ with the minimum number of trees. Therefore, $\mathcal{F}$ is a maximally connected partition of $B$. □

Using Proposition 5.1, we perform a DFS traversal and partition $V$ into $V^+$ and $V^-$ such that the number of inversion edges is minimized, meaning that during DFS traversal, any switch in orientation increases the number of inversion edges.

### 6 Cycle Spaces and Variant Edges

A choice of reference tree for a pangenome graph is akin to a choice of basis for a vector space, specifically the cycle space of the underlying undirected graph. To fully understand this, it is necessary to introduce Eulerian subgraphs, which are subgraphs where each vertex has an even degree. The cycle space of a graph is a vector space formed by cycles over the field of two elements, using symmetric difference as the operation. The symmetric difference of two spanning subgraphs is the subgraph with edges that appear in exactly one of the original subgraphs. The fundamental basis of these cycle spaces consists of cycles formed by adding edges to a spanning tree, providing a minimal set of cycles that can generate any Eulerian subgraph through symmetric differences.

Considering a bidirected graph $B$, the underlying undirected representation, denoted $U(B)$, is a multigraph where each node corresponds to a binode in $B$, and nodes $u$ and $v$ are adjacent if $\{u, u^c\}$ and $\{v, v^c\}$ are adjacent in $B$.

From this foundational setup, the following sequence of propositions proves that any genetic variant (i.e. symmetric differences of given walks) can be captured using variant edges:

**Proposition 6.1.** *The reference tree $T$ of a bidirected graph $B$ induces a spanning tree of $U(B)$, denoted by $U(T)$. Any variant edge in $B$ induces an edge in $U(B)$ that is not an edge of $U(T)$.*

*Proof.* By the definition of the underlying undirected graph, $U(T)$ is connected. Suppose $U(T)$ contains a cycle called $C$. Since $T$ has no edge from any node $u$ to $u^c$, $C$ is not a loop (cycle with one node). Therefore, we may assume that $C$ contains at least two nodes called $u, v$ and hence there are two internally disjoint paths, between $u$ and $v$. By the definition of the underlying undirected graph these two internally disjoint paths induce two paths from $(u, u^c)$ to $(v, v^c)$ in $T$ that contradicts the definition of the reference tree (see section 2). □

**Proposition 6.2.** *The symmetric difference of any two walks in $B$ that start and end at terminus nodes induces an Eulerian subgraph in $U(B)$.*

*Proof.* The degrees on the internal nodes are even in the symmetric difference since for each enter of a walk to a node it must exit once.

For the start terminus nodes each walk exits once and for the end terminus nodes each walk enters once. Therefore, in the symmetric difference the degree of the terminus nodes are either 0 or 2 depending on weather the two walks follow the same edge to exit or inter the terminus node or not. □

**Proposition 6.3.** *Every variant edge in $B$ induces a cycle in $U(B)$. The set of these cycles forms the fundamental basis of the cycle space of $U(B)$.*

*Proof.* Suppose $T$ is a reference tree of $B$. Using Proposition 6.1, $U(T)$ is a spanning tree of $U(B)$ and any variant edge is not an edge of $U(T)$. Therefore, as established in Theorem 6.3.1 of West's textbook [2], the fundamental cycles obtained by adding edges to a spanning tree form a basis for the cycle space of a connected graph.

□

### 7 Reference and Alternative Alleles

Recall that the half-trees of a bidirected tree $T = (V, E, C)$ are the directed subgraphs $T^+ = (V^+, E^+)$ and $T^- = (V^-, E^-)$. $T^+$ is rooted. Let $e = (u, v)$ be a non-tree edge; in particular, it is allowed to be an inversion.

Given a variant edge $(u, v)$ in the bidirected pangenome graph $B$ and reference tree $T$, the *branch point* is defined as the lowest common ancestor of $u^+$ and $v^+$ in the half-tree $T^+$. The complement of the branch point is highest common descendant of $u^-, v^-$ in $T^-$. Intuitively, the branch point marks the divergence between the reference path and the alternative path corresponding to the variant. It is the deepest node in $T$ from which both $u$ and $v$ are reachable via directed paths. The branch point is used to compute reference and alternative alleles and is central to variant annotation, repeat detection (see section 9), and graph simplification (see section 11).

Let $T'$ be the half-tree containing $u$, and let $v' = v$ or $v^c$, whichever one is contained in $T'$. Let $b$ be the lowest common ancestor or highest common descendant of $u, v'$ in $T'$ (i.e., the branch point or its complement). Add $(u, v)$ to $T'$. If $v \neq v'$, also add $v$, and also add the undirected edge $(v', v)$.

In this graph, there exists one pair of parallel paths: one of the endpoints is $b$, and the other is either $u$ or $v$. Call the latter endpoint $a$. One of those two paths contains the member of $u, v$ which is not $a$; this is the "alternative path", and the other is the "reference path". Figure 3 shows reference and alternative paths for different scenarios.

The allele corresponding to a directed path $u_1, ..., u_n$ is the concatenation of the sequences of the nodes, except that when the path visits $u$ and $u^c$ consecutively, these sequences are skipped. Visually, this corresponds to a path that enters and exits from the same 'side' of a binode (Figure 4).

**Proposition 7.1.** *For a variant edge $e = (u, v)$:*

- *If $u$ and $v$ are both positive or negative, then $Ref(e^c) = (Ref(e))^c$ and $Alt(e^c) = (Alt(e))^c$.*
- *If $u$ and $v$ have different orientations, then $Ref(e) = Alt(e^c)$ and $Alt(e) = Ref(e^c)$.*

*Proof.* Let $e = (u, v)$ be a variant edge and let $e^c = (v^c, u^c)$ denote its complement. Let $b$ denote the branch point of $e$, and let the reference and alternative paths be defined as in the caption of Figure 3.

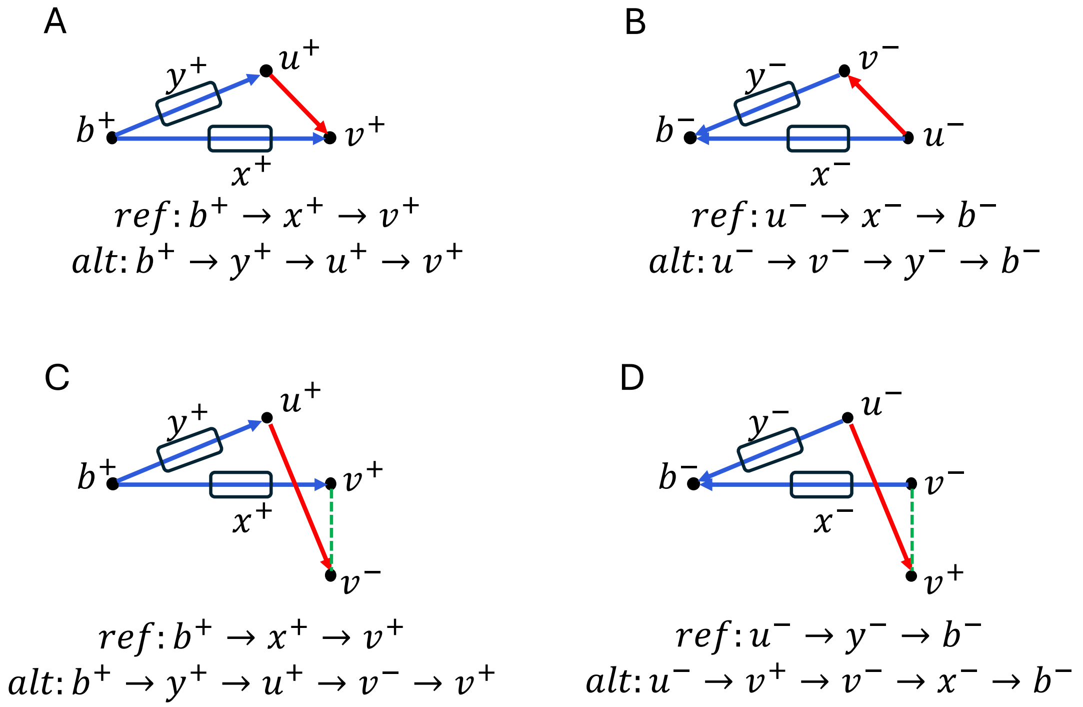


Figure 3: **Reference and alternative paths for all orientations of a variant edge.** Each panel shows the reference path and the alternative path for a variant edge (red). Edges in the reference tree are shown in blue. Panels (A) and (B) illustrate cases where both endpoints of the variant edge lie in the same orientation: (A) forward ($u^+ \to v^+$) and (B) reverse ($u^- \to v^-$). In these cases, the branch point and both endpoints of the variant edge lie within the same half-tree. The two internally disjoint paths from the branch point to the variant endpoint form the reference and alternative paths, distinguished by whether or not they include the variant edge. Panels (C) and (D) illustrate inversion edges: (C) forward to reverse ($u^+ \to v^-$), and (D) reverse to forward ($u^- \to v^+$). In these cases, the second node of the variant edge lies in the opposite half-tree from the branch point and the first node. The green dashed undirected edge represents a switch within the binode that allows traversal between orientations. Reference and alternative paths remain a pair of internally disjoint parallel paths; the alternative path traverses both the variant edge and the green switch edge.

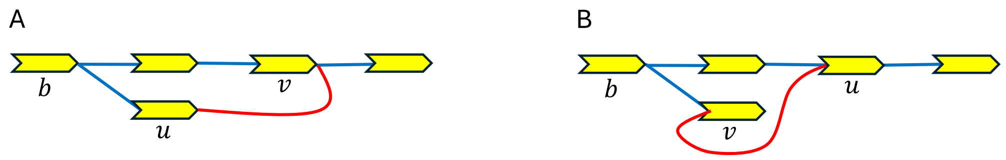


Figure 4: **Illustration of allele simplification when a path visits both orientations of the same binode**. In both panels, the reference path (blue) follows the linear reference sequence. (A) The alternative path traverses nodes $v$ and $u^c$ but does not include both $u$ and $u^c$ or both $v$ and $v^c$ consecutively, so no cancellation occurs, and the alternative allele includes the sequence of $u^c$. This is visually clear from the figure since the alternative path enters and exists from different sides of binodes $u$ and $v$ when traversin the variant edge. (B) The alternative path contain both $u, u^c$ of $v, v^c$ and This results in a simplified allele with the corresponding segment omitted. This is visually clear since the alternative path enters and exits the same binode from the same side (either $u \to u^c$ or $v \to v^c$ depending on the orientation of the treversal).

**Case 1: $u$ and $v$ have the same orientation.** Then $e$ and $e^c$ lie entirely within the same half-tree (either $T^+$ or $T^-$), and the reference and alternative paths of $e^c$ are exactly the reverse complements of those of $e$. Therefore, $\text{Ref}(e^c) = (\text{Ref}(e))^c$ and $\text{Alt}(e^c) = (\text{Alt}(e))^c$.

**Case 2: $u$ and $v$ have different orientations.** In this case, $e$ is an inversion edge, and its complement $e^c$ traverses the same binodes in reverse order but with reversed strand orientation. From Figure 3, we observe that the alternative path of $e$ corresponds to the reference path of $e^c$, and vice versa, since switching the direction reverses the role of the tree-consistent and variant-containing paths. Therefore, $\text{Ref}(e) = \text{Alt}(e^c)$ and $\text{Alt}(e) = \text{Ref}(e^c)$. □

**Efficient computation of reference and alternative alleles:** To efficiently compute reference and alternative alleles, we need to efficiently compute branch points for all variant edges. We use *Tarjan's offline lowest common ancestor algorithm* [1]. This classic algorithm performs a single linear-time traversal of the tree and answers each LCA query in nearly constant time, making it well-suited for large pangenome graphs with many variants.

### 8 Variant Types

Variant edges in the pangenome graph are classified into three reference-dependent categories based on their structural characteristics relative to the reference tree. These categories help in understanding the genetic variations and their implications:

- **Forward Edges:** These correspond to deletions, where the walk skips over a segment that is present in the reference tree. Such edges imply that a segment found in the reference genome is absent in the alternate genome.
- **Backward Edges:** These represent duplications, where the walk traverses a segment of the genome multiple times. This implies that a section of the genome is duplicated in the alternate path compared to the reference.

- **Crossing Edges:** These signify replacements, where one segment of a walk is replaced by another segment. Single Nucleotide Polymorphisms (SNPs) and Multiple Nucleotide Polymorphisms (MNPs) are replacements whose reference and alternative alleles are both one basepair long and both $n$ basebairs long, respectively.
- **Inversions:** These edges connect nodes with opposite orientations (positive and negative directions) and result in a segment of the genome being reversed in orientation relative to the reference sequence. An inversion event corresponds to a *pair* of such inversion edges: the first edge switches the walk from forward to reverse orientation, and the second brings it back to the forward direction (or vice versa). Only when these inversion edges are properly paired can the walk rejoin the reference strand at the correct orientation; otherwise, the walk remains stranded in the reverse direction and cannot complete a valid traversal from one end of the genome to the other.

### 9 Repeats

The pangenome graph utilized in our study is based on the minigraph-cactus framework. Due to the construction methodology of minigraph, the resulting graph structure is almost a Directed Acyclic Graph (DAG), with very few back edges with respect to the DFS reference tree. This characteristic significantly reduces the occurrence of repeat variants within the graph. However, the node sequences themselves often contain repeated motifs. To address this, we developed and implemented an algorithm specifically designed to identify such repeat sequences within the nodes of the pangenome graph.

**Algorithm 9.1.** *The algorithm identifies repeat motifs for a given variant edge:*

1. *Ensure the variant edge is not an inversion.*
2. ***Select Allele:*** *Choose the reference or alternate allele, depending on which is non-empty.*
3. ***Detect Motif:*** *For each divisor length $r$ of the allele's length, check if the allele can be written as a perfect repeat of a substring of length $r$.*
4. ***Verify in Graph Context:*** *Check for the motif upstream of the branch point or downstream of the variant target node. If found, annotate it.*

### 10 Missingness

In practice, haplotype assemblies are not fully contiguous; they contain gaps, due to the difficulty of assembling repetitive regions, and are represented as collections of walks. Between the end of one walk and the beginning of the next, there may exist many possible paths through the graph. We define a variant edge as being missing for a haplotype if that edge could be visited in between the end of one walk and the beginning of the next one. Directly identifying missing edges is computationally challenging, requiring a graph traversal for every walk. However, variant positions can be used to identify most missing edges.

Edges that violate the topological order inherited from the reference tree, when restricted to positive nodes, are called back edges. Removing the back edges from the graph restricted to positive nodes results in a directed acyclic graph (DAG). The right position of a positive node $u^+$ is defined as the smallest position of any node on the linear reference that is reachable from $u$ in the DAG. Similarly, the right position of a binode $(u^+, u^-)$ is defined as the right position of $u^+$ and denoted by $\text{rpos}(u)$.

For a walk $W = ((u_1^+, u_1^-), \ldots, (u_k^+, u_k^-))$, the linear coverage of $W$ is the interval $[a, b]$ where $a := \min_{u \in W}\{\text{pos}(u)\}$ and $b := \max_{u \in W}\{\text{rpos}(u)\}$. We identify as missing those variants of a haplotype to be the variant edges whose positions do not intersect the union of linear coverages of walks belonging to that haplotype. This approach identifies as missing all variant edges which can be traversed by a path that begins at the endpoint of one walk, terminates at the start point of the next walk, and visits some node on the linear reference (which could be one of those endpoints). However, it should be noted that this method may fail to detect a missing variant edge $(u, v)$ that could be visited between the end of one walk, $W_1$, and the beginning of the next walk, $W_2$, if the right position of $v$ is greater than $\min_{u \in W_2}\{\text{pos}(u)\}$. This situation arises if the beginning of $W_2$ is reachable from $v$ via a back edge $(a, b)$ with $\text{pos}(a) > \min_{u \in W_2}\{\text{pos}(u)\}$ and $\text{pos}(b) \leq \min_{u \in W_2}\{\text{pos}(u)\}$.

Algorithm 10.1 identifies missing variants for each haplotype.

**Algorithm 10.1.** *The algorithm identifies missing variants for each haplotype using the positions and coverage of its walks:*

1. ***Compute Walk Coverage:*** *Compute the linear coverage interval $[a_W, b_W]$ where:*

$$a_W := \min_{u \in W}\{pos(u)\}, \quad b_W := \max_{u \in W}\{rpos(u)\}$$

2. ***Aggregate Haplotype Coverage:*** *Sum edge counts across walks and merge intervals to get total coverage for the haplotype.*
3. ***Detect Missing Edges:*** *Mark any unvisited variant edge as missing if its position does not intersect the haplotype's coverage.*

### 11 Simplified Graph

In complex regions of the pangenome graph, such as bubbles with multiple variant edges, numerous small variants often lies within larger variant structures, such as long indels with several SNPs on them. To simplify the analysis and visualization of these complex regions, we designed and implemented an algorithm that reduces the graph to a minor that retains only the long variants. This simplification helps in distilling the essential topological features of complex genomic regions.

**Algorithm 11.1.** *The algorithm simplifies the subgraph of a complex genomic region by removing small variants:*

1. ***Filter by Allele Length:*** *Remove variant edges whose total allele length is below a given threshold.*
2. ***Identify Endpoints:*** *If endpoints are not provided, identify all nodes on the reference path with either in-degree or out-degree equal to zero.*
3. ***Delete Tips:*** *Iteratively remove tip-like structures that do not lie on paths between identified endpoints.*
4. ***Contract Simple Paths:*** *Merge linear paths—that is, paths in which all internal nodes have both in-degree and out-degree equal to one—between endpoints.*

#### 12 Handling Non-Path Linear References

In some pangenome graphs, such as those produced by PGGB, the linear reference genome (e.g., GRCh38 or CHM13) is not embedded as a simple path. Rather, it may be represented as a walk that visits certain nodes or edges multiple times. The linear reference has a special significance in our approach because we use it to assign positions; however, we can still construct a valid reference tree and assign positions to all nodes and variant edges in a principled way.

##### 12.1 Reference Tree Construction

Let $L$ denote the linear reference walk in the bidirected graph $B$. Suppose the binodes of $B$ have already been oriented as described in Section 5.

Since we assume the bidirected graph has already been oriented, if the reference walk $L$ contains any inversion edges (edges between nodes of opposite orientation), we discard those edges from $L$ before constructing the reference tree. The remaining edges of $L$ are then used to construct a spanning DFS tree $T_L = T_L^+ \cup T_L^-$. This tree includes all nodes visited in the forward and reverse orientations along the walk and consists of two disjoint half-trees: one rooted DFS tree on $V(L)^+$ (denoted $T_L^+$) and one directed tree on $V(L)^-$ (denoted $T_L^-$).

Next, we extend this DFS tree into a full spanning tree $T = T^+ \cup T^-$ of the entire graph $B$ using a depth-first traversal starting from $T_L^+$. This process generalizes our construction for path-based linear references, where the linear reference path formed the first branch of the traversal. Here, the traversal begins from a rooted DFS tree, $T_L$. The resulting spanning tree $T$ remains a valid reference tree for $B$, and all results established for path-based references, such as Theorem 2.1 (that genotypes determine walks up to cycles), and propositions in section 6 continue to hold.

##### 12.2 Position Assignment

When the linear reference $L$ is not a path but a walk, certain nodes may be visited multiple times. This introduces ambiguity in defining a unique position for such nodes. To address this, we define positions as follows.

Let $u \in V$ be a node not on the linear reference walk $L$. Define $\text{pos}(u)$ to be the position of its lowest ancestor in $T_L$. If $u$ lies on $L$ and is visited multiple times, we associate it with a set of positions (one for each occurrence in the walk) then assign $pos(u)$ to be the set of all such positions.

For a variant edge $e = (u, v)$, we define its position $pos(e)$ to be the smallest interval that contains all positions of $u$ and $v$.

This position definition retains key theoretical guarantees. In particular, an analog of Theorem 4.1 holds: the genotype count $x_e(w)$ for any variant edge $e$ in a walk $w$ is determined by the counts of those variant edges whose positions overlap $\text{pos}(e)$. Similarly, variant edge positions remain consistent under complementation, and variant reconstruction remains valid.

We still prefer to define positions with respect to a linear reference path. When the reference walk contains long cycles, it causes faraway positions to be superimposed, effectively making it useless to define any genomic region smaller than the length of the cycle. This is particularly suboptimal for recent SNPs that occurred after the emergence of a duplication event: such SNPs naturally belong to one copy or the other, not equally to both of them.
