## Supplementary Figures for "Defining and cataloging variants in pangenome graphs"

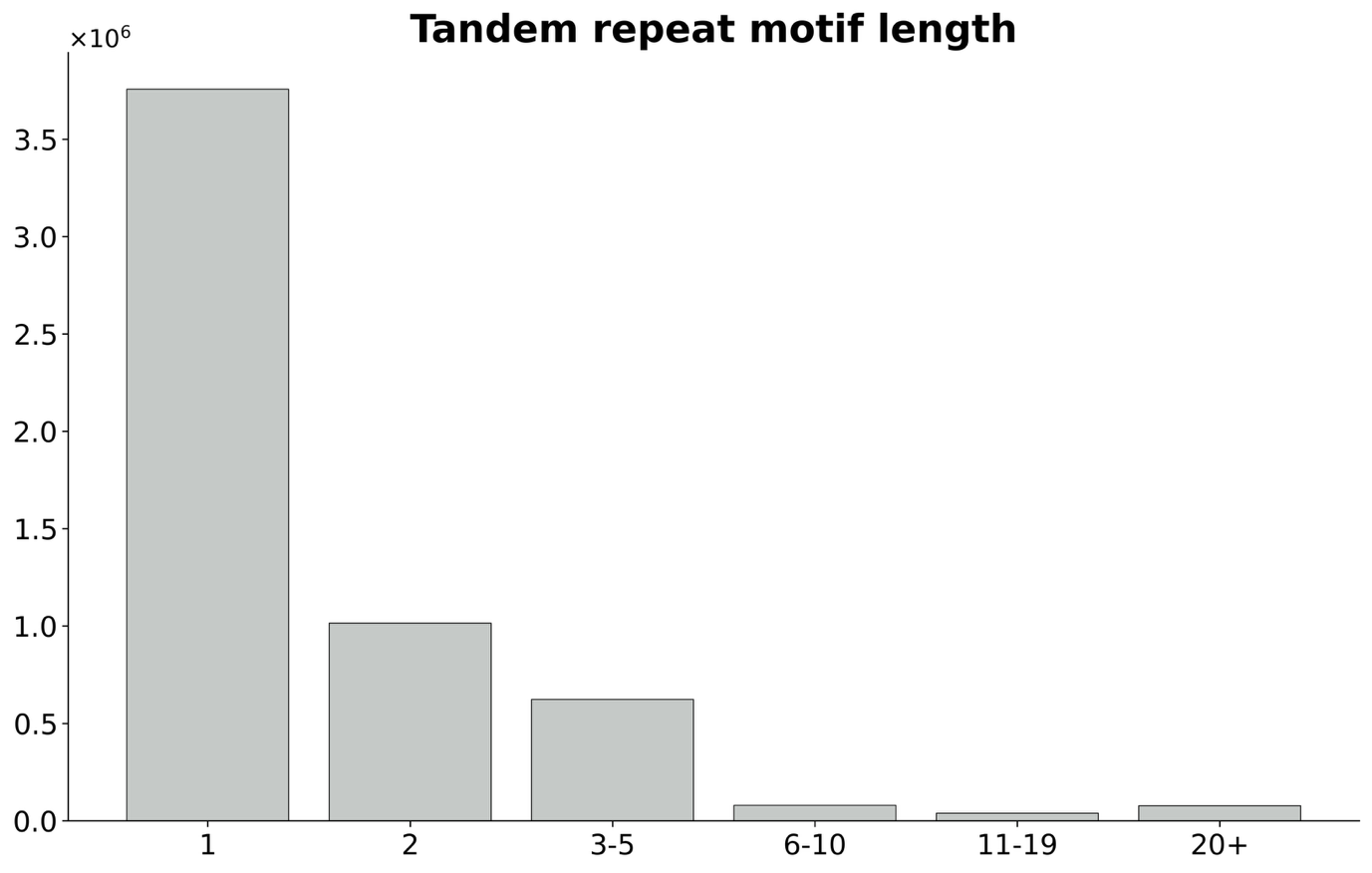
**Supplementary Figure 1: distribution of tandem repeat motif lengths.** Tandem repeat motif length was defined as the length of the length of the ‘TR_motif’ field in the pantree VCF.

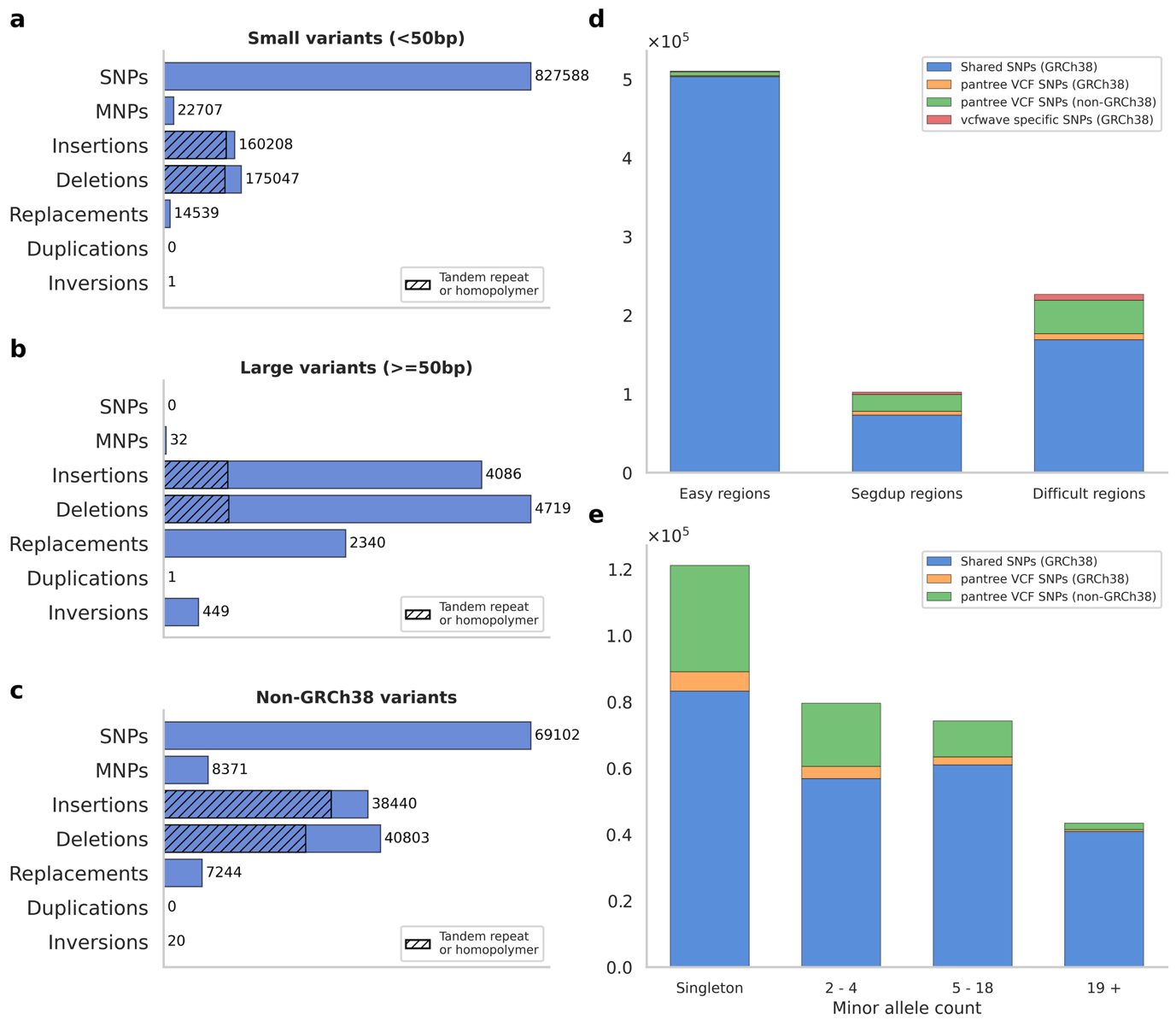
**Supplementary Figure 2: variants on chromosome X.** (a) Number of small variants (<50bp combined allele length) across seven non-overlapping variant types. Hatched area indicates the number of variants that cause a change in the copy number of a reference-tree basepair sequence. (b) Number of large variants. (c) Number of non-GRCh38 variants, defined as variant edges connecting two non-GRCh38 nodes. (d) Overlap of SNPs with those reported by *vcfwave*, across three non-overlapping genome annotations. (e) Minor allele count (the minimum of the reference allele count and alternative allele counts) out of 90, stratified by overlap with *vcfwave*.

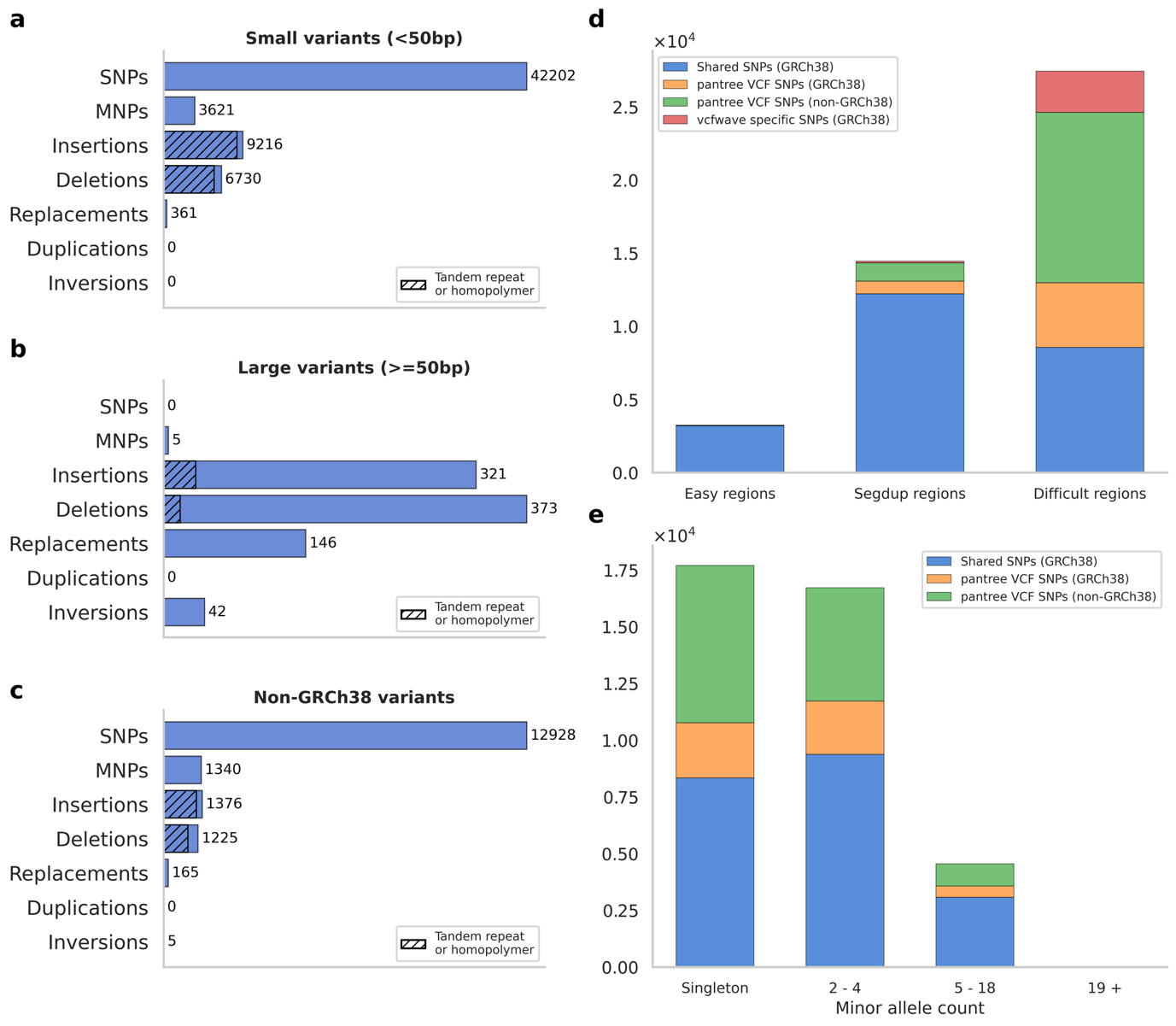
**Supplementary Figure 3: variants on chromosome Y.** (a) Number of small variants (<50bp combined allele length) across seven non-overlapping variant types. Hatched area indicates the number of variants that cause a change in the copy number of a reference-tree basepair sequence. (b) Number of large variants. (c) Number of non-GRCh38 variants, defined as variant edges connecting two non-GRCh38 nodes. (d) Overlap of SNPs with those reported by *vcfwave*, across three non-overlapping genome annotations. (e) Minor allele count (the minimum of the reference allele count and alternative allele counts) out of 21, stratified by overlap with *vcfwave*.

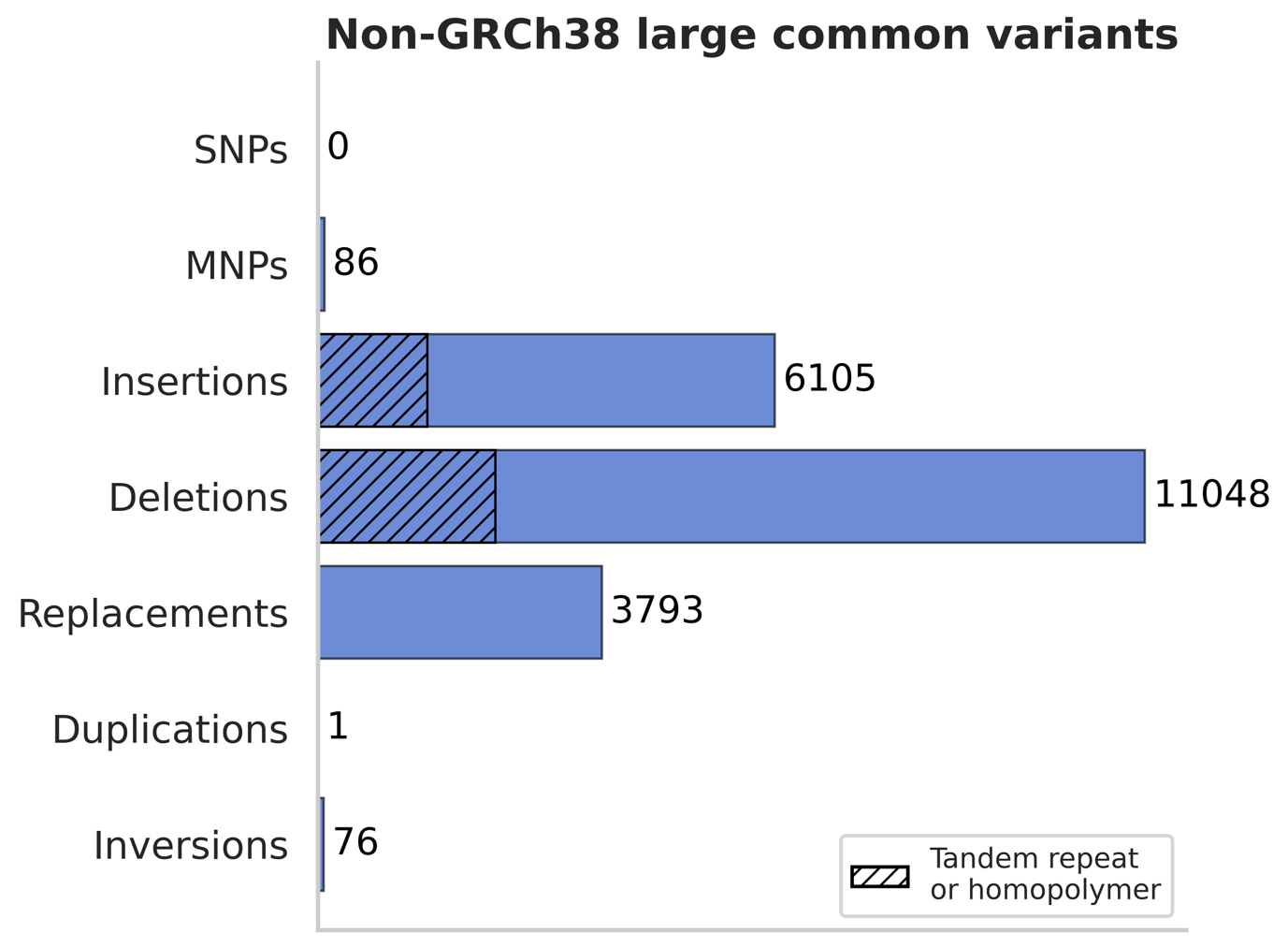

**Supplementary Figure 4: large, common non-GRCh38 variants.** Total counts per variant class (SNP, MNP, insertion, deletion, replacement, duplication, inversion) for non-GRCh38 variants with a minor allele count of at least 5 and a combined allele length of at least 50bp. Hatched bars denote variants that cause a repeat expansion or contraction, including all duplications.

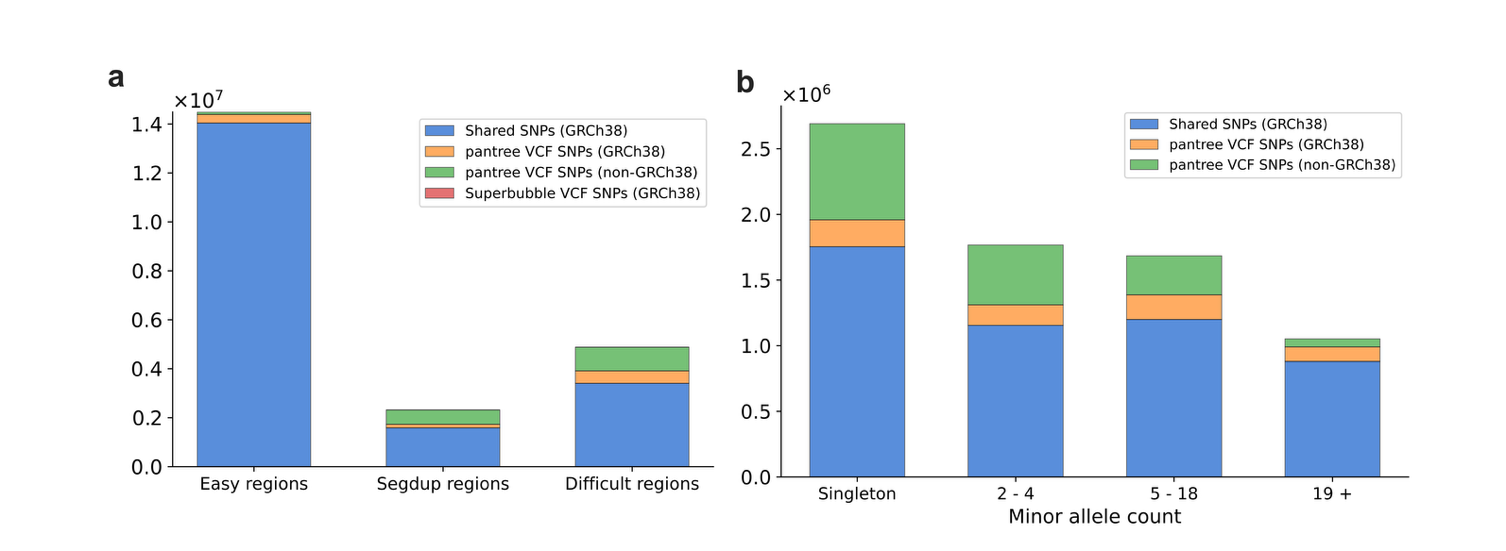

**Supplementary Figure 5**: SNPs detected using pantree vs. superbubbles. (a) SNPs were partitioned by their VCF position into easy regions, segmental duplications, and other difficult regions. Bars indicate the number of SNPs detected using both methods, the number of GRCh38 and non-GRCh38 SNPs found by pantree only, and the number of SNPs found using superbubbles only (there were exactly 2 of these). (b) SNPs separated by their minor allele count, defined using pantree (the minimum value between reference allele counts and alternative allele count). Superbubble VCF specific SNPs were not included because they did not have allele counts reported.

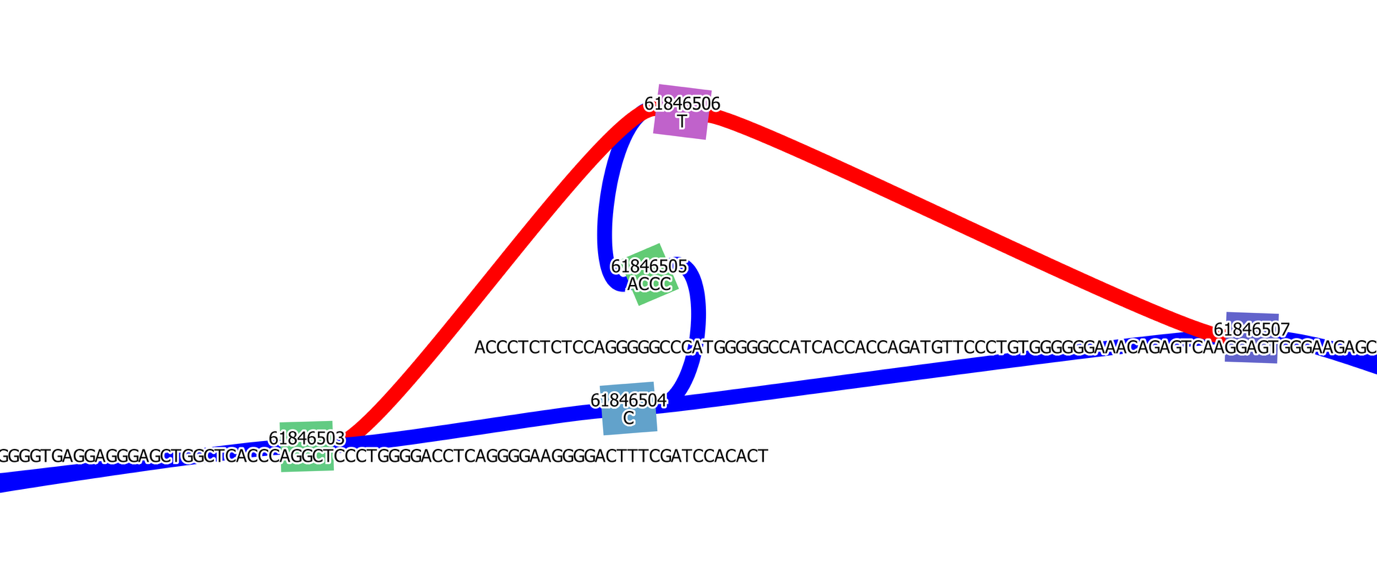
**Supplementary Figure 6: a superbubble-specific SNP.** Exactly two SNPs were detected as superbubbles without being called as SNPs by pantree. This superbubble, on chromosome 6, coincides with an assembly gap; there exists a terminal edge from the negative starting terminus to the node 61846506, which means that one walk starts in the middle of the superbubble from the node 61846506, then goes through the nodes 6184605 and 6184604, then to the pinch-point 6184603. Because superbubble alleles are required to pass through both ends of the superbubble, this walk, whose allele would be ’CACCCT’, is not counted as an alternative allele, and the superbubble is classified as a SNP. pantree identifies an insertion (‘ACCCT’) and a deletion (‘ACCC’).

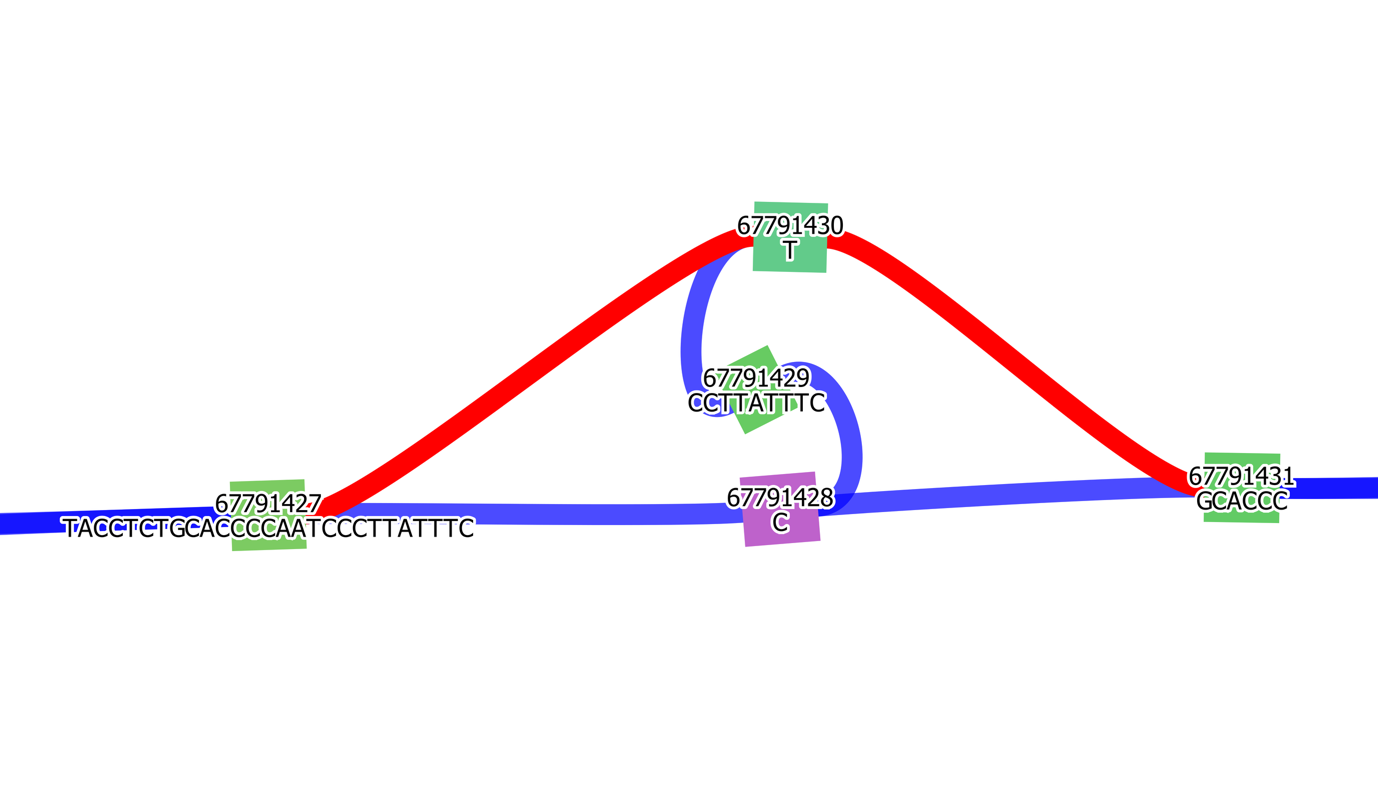
**Supplementary Figure 7: a superbubble-specific SNP.** Exactly two SNPs were detected as superbubbles without being called as SNPs by pantree. This superbubble, on chromosome 7, coincides with an assembly gap (similar to the SNP in Supplementary Figure 6). There is a terminal edge from the node 67791428 to the negative ending terminus, and a walk which begins within node 67791431 following the negative orientation, then goes through the nodes 67791430 and 67791429, then ends at 67791428. Thus, the ’CCCTTATTTCT’ allele did not show up as an alternative allele in this superbubble.

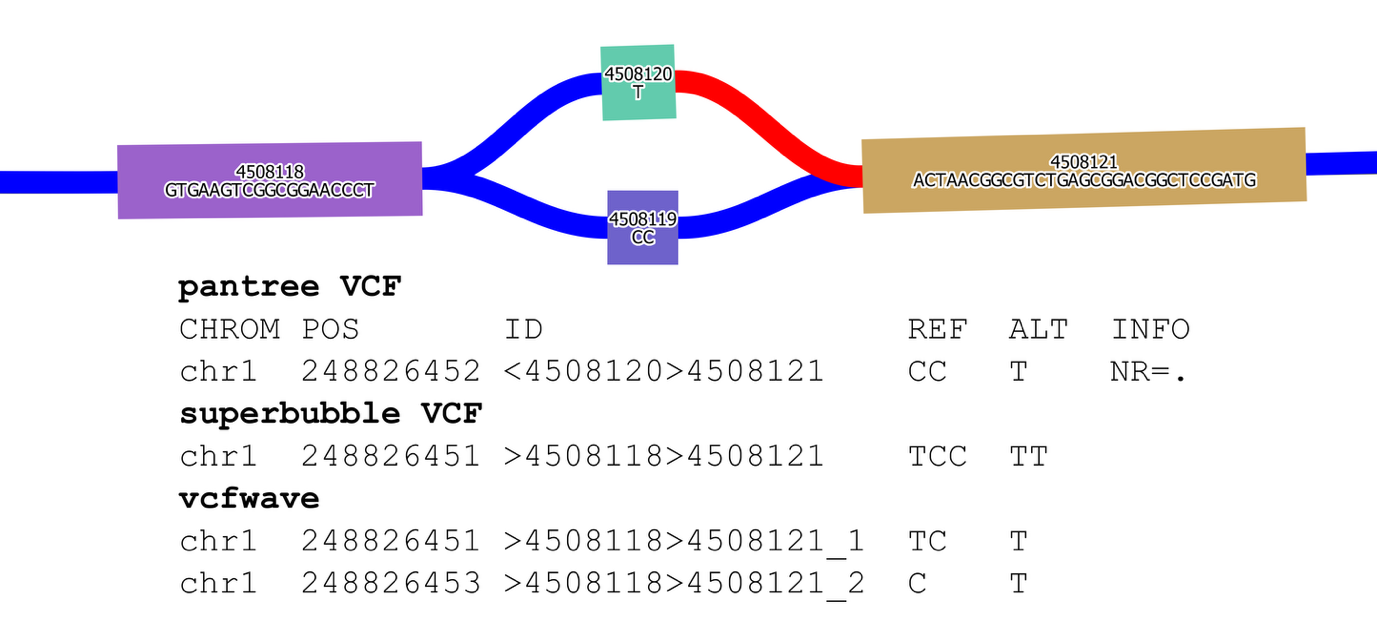

**Supplementary Figure 8: a *vcfwave*-specific SNP that is inconsistent with the graph.** pantree identifies a replacement whose reference allele is CC and alternative allele is T, and the ‘raw’ superbubble VCF has a similar variant (except prepending the last letter of the branch point to both alleles). *vcfwave* produced a different alignment and identified two variants, including a C-to-T SNP which is inconsistent with the graph.

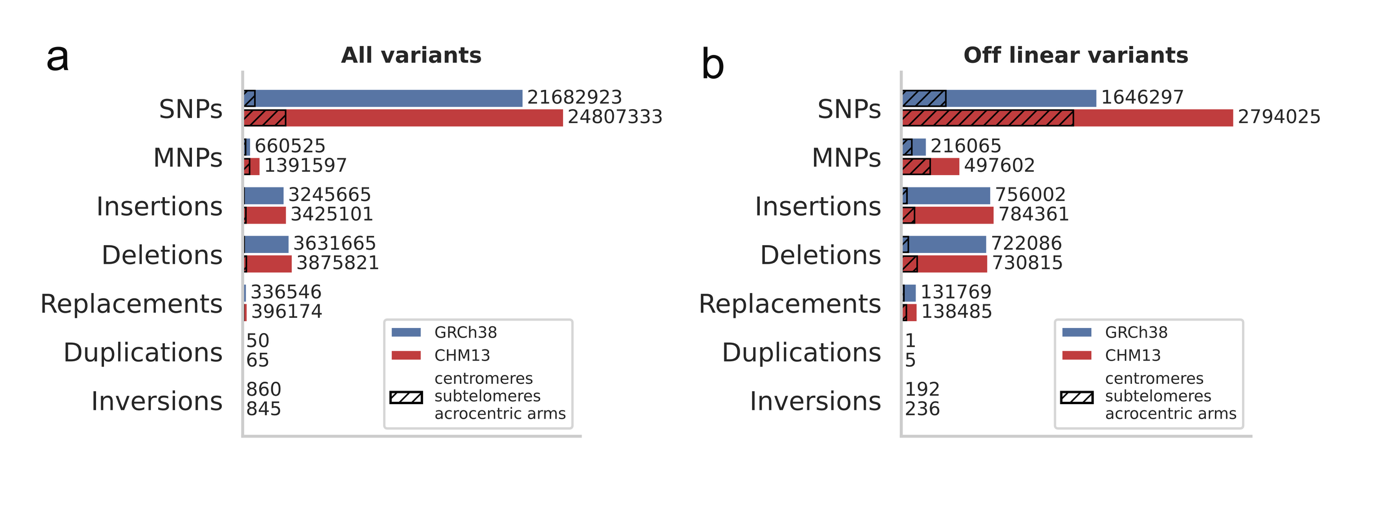
**Supplementary Figure 9: variant comparison between the CHM13 and GRCh38 graphs.** These graphs are based on the same data and the same algorithm, Minigraph-Cactus, but the algorithm is initialized at one or the other linear reference genome. The graphs differ in combined sequence length because regions missing from GRCh38 are excluded from the GRCh38 graph. These non-GRCh38 regions mostly include centromeres, subtelomeres, and acrocentric chromosome arms. (a) The number of variants detected by pantree in each graph. (b) The number of non-GRCh38 variants in the GRCh38 graph and the number of non-CHM13 variants in the CHM13 graph.

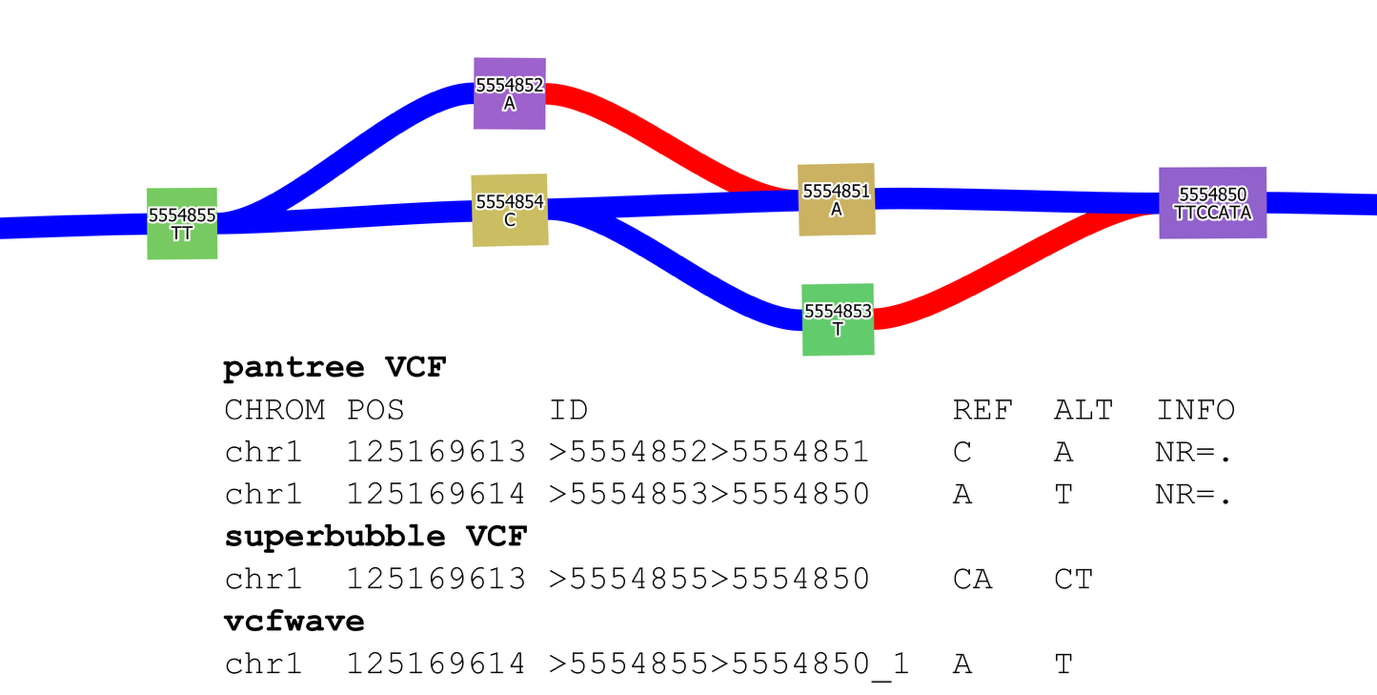
**Supplementary Figure 10: a SNP missed by *vcfwave* due to an assembly gap.** pantree identifies two SNPs. One walk terminates at the node 5554851, and as a result, the ‘AA’ allele is not listed in the superbubble ‘raw’ VCF. As a result, the *vcfwave* VCF misses the ‘C’ to ‘A’ SNP.

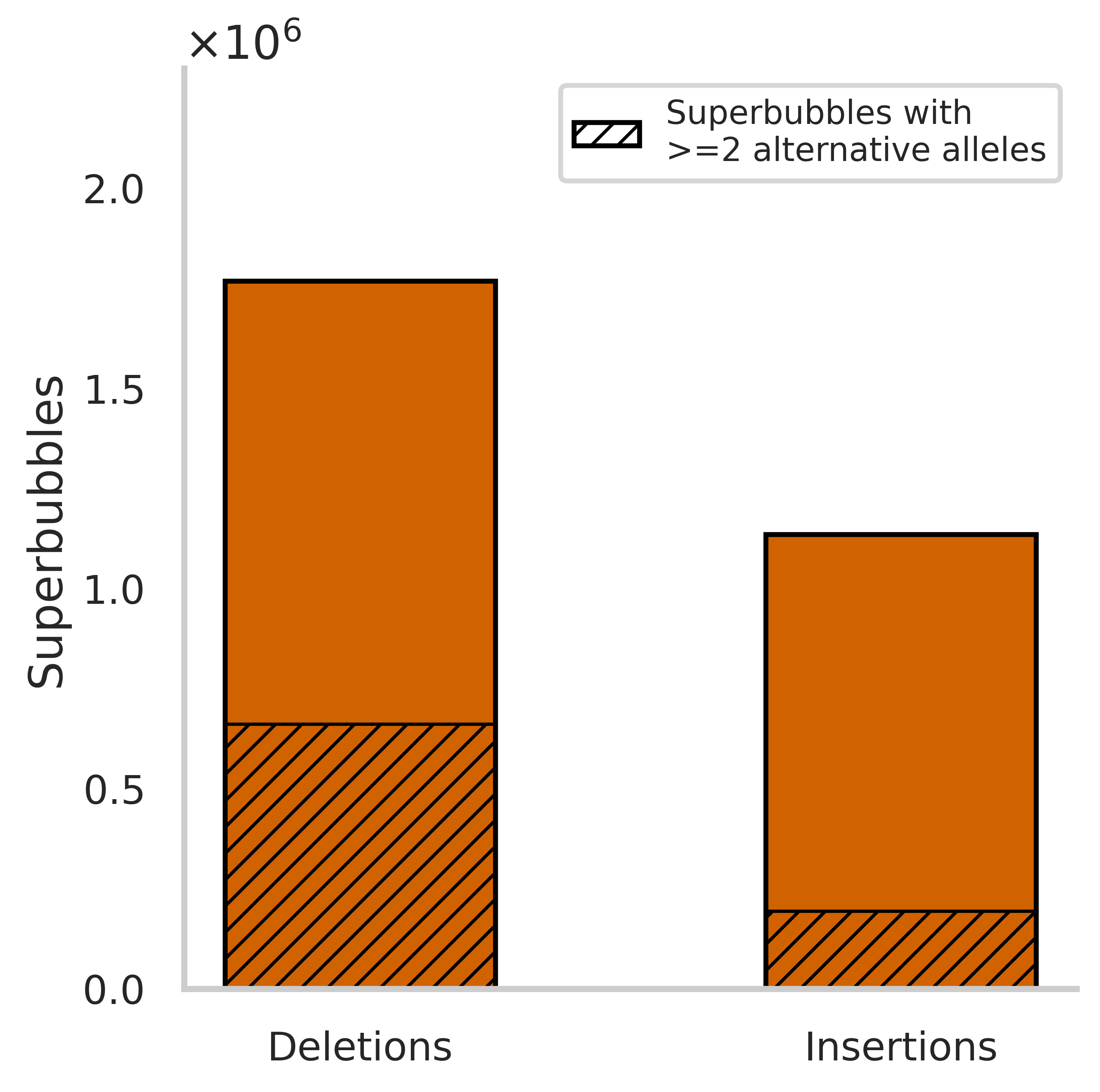

**Supplementary Figure 11: Proportion of insertion and deletion superbubbles.** Among all superbubbles, there are 55.7% more deletion superbubbles (1,767,792) than insertion superbubbles (1,135,061). Among all multiallelic superbubbles, there are 341.6% more deletion superbubbles (661,196) than insertion superbubbles (193,553). 37.4% of deletion superbubbles are multiallelic, versus 17.1% of insertion superbubbles.
