## Supplementary Tables List for "Defining and cataloging variants in pangenome graphs"

**Supplementary Table 1: Summary of variants identified.** This table summarizes variants identified across chromosomes 1-22, used to generate Figures 2a-c. The first column denotes the chromosome, with “all” indicating the total across the 22 chromosomes. The second column specifies variant types (SNP, MNP, insertion, deletion, replacement, inversion, duplication), categorized by allele length (small/large), minor allele count (common/uncommon), and reference status (GRCh38/non-GRCh38).

**Supplementary Table 2: Summary table of the tandem repeats.** This table is used to mark the hatched part of the Figure 2a, 2b, and 2c. It has the same structure as Supplementary Table 1, but each entry specifies the number of tandem repeats/homopolymers.

**Supplementary Table 3:** **Variants summary in different regions.** This table summarizes variants within easy regions, segmental duplications and hard regions of chromosomes 1-22.

**Supplementary Table 4: Comparison of SNPs found by different methods.** This table is used to generate Figure 2d. It compares the number of SNPs identified by pantree (‘ourvcf’), *vcfwave*, and the superbubble approach (‘rawvcf’) on chromosomes 1-22, stratified by region. ‘Shared’ indicates the number of SNPs shared between pantree and the second method; ‘Ourvcf_only’ indicates the number of SNPs found by pantree but not the second method; ‘Vcfwave_only’ and ‘Rawvcf_only’ indicate the number of SNPs found by the second method but not pantree. ‘linear’ and ‘offlinear’ indicate the number of GRCH38 and non-GRCh38 variants, respectively.

**Supplementary Table 5: Frequency-stratified comparison of SNPs found by different methods.**  This table is used to generate Figure 2e. It compares the number of variants in segment duplications and difficult regions identified by each method at different allele count strata: 1, 2-4, 5-18, and 19+ out of 90. For column descriptions, see Supplementary Table 4 caption.

**Supplementary Table 6: Distribution of variant edge count and alternative allele count across superbubbles.** This table is used to generate Figure 3b. Superbubbles are binned by their number of variant edges or alternative alleles, on chromosomes 1-22.

**Supplementary Table 7: Superbubble frequency by variant and ALT counts.** This table is used to generate Figure 3d. It summarizes the number of superbubbles observed for each possible combination of variant count and ALT allele count on chromosomes 1-22.

**Supplementary Table 8: Counts of triallelic superbubbles by type.** This table is used to plot Figure 3f. It summarizes the count distribution of triallelic superbubble types on chromosomes 1-22.

**Supplementary Table 9: Variant summary in triallelic superbubbles.** This table summarizes the variants contained in superbubbles with 2 variant edges (triallelic), stratified small vs. large and GRCh38 vs. non-GRCh38, on chromosomes 1-22.

**Supplementary Table 10: Summary of superbubbles by type, ALT Allele group, and variant counts.** This table is used to plot Figure 3c and Figure 3g-h. It bins superbubbles on chromosomes 1-22 by ALT allele count group and stratifies them as insertions, deletions or neither. It counts the number of superbubbles, the number of superbubbles containing at least one tandem repeat (TR)-annotated variant, and the number of variants contained in those superbubbles; variants are stratified TR vs. non-TR and GRCh38 vs. non-GRCh38.

**Supplementary Table 11: Genomic range of HLA and RHD.** This table compares the UCSC-defined position ranges for the HLA and RHD regions with the graph-derived ranges, which are determined by selected start and end nodes surrounding each region.

**Supplementary Table 12: Summary of highly allelic superbubbles with long insertions.** This table presents the counts of superbubbles with at least 10 alternative (ALT) alleles. A subset of these superbubbles also contains insertions of at least 1kb. For each category, the number of total superbubbles, as well as the counts of GRCh38 and non-GRCh38 variant edges, are reported.
